# MALDI-TOF imaging mass spectrometry demonstrates sex- and age-dependent spatial changes in brain energy metabolism in response to amyloid stress using a mouse model of Alzheimer’s Disease

**DOI:** 10.64898/2026.09.14.750669

**Authors:** Sandra Grahovac-Nemeth, Kristina Jurcic, Marc Courchesne, Karen Nygard, Grace Callahan, Ariel K. Frame, Reza Khazaee, Wenxuan Wang, Moganatharsa Ganeshalingam, Raymond Thomas, Shawn N. Whitehead, Robert C. Cumming

**Affiliations:** Department of Biology, Western University, London, ON, N6A 5B7, Canada; MALDI Mass Spectrometry Facility, Western University, London, ON, N6G 2V4, Canada; Biotron Integrated Microscopy Facility, Western University, London, ON, N6A 3K7, Canada; Department of Anatomy & Cell Biology, Western University, London, ON, N6A 5C1, Canada

**Keywords:** AD – Alzheimer’s disease, Aβ – amyloid beta, AG – aerobic glycolysis, DAN – 1, 5 – diaminonaphthalene, lactate, lactylation, MALDI-TOF imaging, metabolites, APP/PS1 mice

## Abstract

Alzheimer’s disease (AD) is the most common form of dementia, and no therapies currently exist that prevent or slow its progression. Lactate has recently emerged as both an energy substrate and a signaling molecule required for memory formation, acting in part through a novel epigenetic mechanism termed histone lactylation. Here, we used matrix-assisted laser desorption/ionization time-of-flight (MALDI-TOF) imaging mass spectrometry and immunofluorescence microscopy to spatially map lactate, glutamate, pyruvate, and citrate levels, alongside histone lactylation, in the brains of wild type and AD transgenic mice at 6 and 18 months of age. Lactate and glutamate were highest in young animals and declined with age, while pyruvate showed the inverse pattern. These shifts were most pronounced in females, and pyruvate-to-lactate and pyruvate-to-citrate ratios suggested a progressive, region-specific shift from glycolytic to oxidative metabolism. Sex was a dominant determinant of cerebral metabolite distribution: females maintained consistently higher lactate and glutamate than males at both ages, indicating a sex-specific metabolic phenotype that persists across physiological aging. Elevated lactate levels were paralleled by increased histone lactylation in aged females, particularly within the cortex and CA2/CA3 hippocampal subregion, and in transgenic females lactylation was enriched in putative microglia near amyloid plaques. Lactate and histone lactylation were positively correlated in wild type mice, consistent with a lactate-driven epigenetic mechanism possibly in microglia, but this relationship was weakened or absent in transgenic mice despite elevated plaque-adjacent lactylation, suggesting amyloid pathology decouples metabolic state from epigenetic regulation. These findings identify sex as a major, underappreciated variable shaping brain metabolic-epigenetic coupling during aging and amyloid stress. Together, these results implicate sex-specific lactate metabolism and lactylation signaling as potential contributors to differential AD vulnerability, and underscore the need to incorporate sex as a biological variable in future studies of metabolic-epigenetic mechanisms and therapeutic targeting in AD.

## Introduction

Alzheimer’s disease (AD) is the most common form of dementia, making up approximately 60 to 70% of dementia-related cases worldwide [1]. Age is considered to be one of the strongest risk factors for AD [2,3], highlighting an increasingly important public health concern due to improved life expectancy and a rapidly growing elderly population arising from the “baby boom” generation [4,5]. In addition, AD is nearly two thirds more prevalent in women than men [6,7]. This sex disparity cannot be fully explained by women’s greater longevity alone, and growing evidence points to sex-specific differences in brain energy metabolism, mitochondrial function, and neuroinflammatory signaling as contributing factors to differential AD risk and progression [8,9]. Despite this, sex is frequently underrepresented as a biological variable in preclinical metabolic studies of AD, leaving the relationship between sex, brain metabolism, and disease vulnerability poorly defined [10].

Amyloid plaques, found extracellularly and in the walls of cerebral blood vessels, are a pathologic hallmark of AD. These plaques contain toxic amyloid beta (Aβ) peptides arising from the cleavage of the amyloid precursor protein (APP) [11,12]. A prevailing theory which has shaped research as well as drug development for over 25 years posits that increased production or inadequate clearance of Aβ are the primary causal events of AD pathogenesis [12,13]. However, recent clinical trials based on removing soluble and/or insoluble Aβ or targeting enzymes responsible for cleaving APP, have demonstrated only modest slowing of cognitive decline and do not halt disease progression [14,15]. Thus, research has moved beyond an amyloid-centric focus and AD is now recognized as a complex and multifaceted disease. To date, over 50 genetic loci have been implicated in AD progression [16]. It is also now recognized that there is a long prodromal phase during which abnormal Aβ is generated at least 25 years prior to clinical evidence of cognitive decline. Furthermore, changes in neuronal function and cerebral metabolism have been shown to precede behavioral abnormalities [17]. Therefore, alternative lines of investigation into related biological pathways and identification of new therapeutic targets are desperately needed to either prevent the development or slow the progression of dementia [18].

Glucose is considered to be the main source of energy in the brain, processed via glycolysis to produce pyruvate, which subsequently fuels mitochondrial oxidative phosphorylation (OXPHOS) [19,20]. However, glucose can also be broken down to generate lactate [21,22]. Interestingly, lactate has historically been considered as either a “waste product” or an indicator of inadequate oxygen supply to a tissue, since it is typically produced when oxygen is rate limiting. The conversion of pyruvate to lactate is catalyzed by the enzyme lactate dehydrogenase A (LDHA), resulting in the net generation of only 2 adenosine triphosphate (ATP) molecules, compared to the potential 32 to 36 ATP that can be produced by mitochondrial OXPHOS [23]. However, cancer cells have been shown to employ an alternative glycolytic metabolism termed “the Warburg Effect” also known as “Aerobic Glycolysis” (AG). Cancer cells break down glucose using AG to predominately generate lactate, even in the presence of sufficient amounts of oxygen, and AG has been shown to occur in the brain as well [24]. Additionally, within the brain, several lines of evidence indicate that AG is cell-type-specific where astrocytes predominantly produce lactate in response to neuronal activity, while neurons preferentially take up astrocyte-derived lactate. This phenomenon forms the basis of the astrocyte-neuron lactate shuttle (ANLS) hypothesis [22,25,26]. Moreover, several studies have shown that astrocyte-to-neuron transfer of lactate operates primarily at excitatory synapses and is necessary for Long Term Potentiation (LTP) as well as memory consolidation [27–29].

This coupling between excitatory neurotransmission and lactate production is thought to be driven in part by glutamate itself: astrocytic uptake of synaptically released glutamate stimulates glucose uptake and glycolysis, thereby increasing local lactate production to meet the metabolic demands of active neurons [25,26]. Beyond its role as a neurotransmitter, glutamate is also tightly linked to cerebral energy metabolism through the glutamate-glutamine cycle, and disruptions in glutamatergic signaling have been implicated in excitotoxicity and synaptic loss in AD [30–33].

In addition to lactate and glutamate, pyruvate and citrate serve as key indicators of the balance between glycolytic and oxidative brain metabolism. Pyruvate sits at the branch point between cytosolic glycolysis and mitochondrial oxidative metabolism, where it may be reduced to lactate by LDHA or transported into mitochondria for entry into the tricarboxylic acid (TCA) cycle [34]. Citrate, an early TCA cycle intermediate, reflects mitochondrial oxidative metabolism while also serving as the principal source of cytosolic acetyl-CoA required for cholesterol and fatty acid synthesis during myelin production [35]. Given that glycolytic and oxidative metabolism are known to shift across the lifespan and may be differentially affected by AD pathology [18], examining pyruvate and citrate alongside lactate and glutamate, and their ratios to one another, offers a more complete picture of metabolic flux than lactate alone.

In a previous study, we observed that cortical lactate levels decrease with normal aging in wild type mice, coupled with an age-dependent decline in AG-related enzymes [36]. In contrast, lactate levels accumulate in the frontal cortex and hippocampus of a transgenic APP/PS1 mouse model of AD. Interestingly, higher expression of lactate-producing enzymes, such as LDHA, correlated with improved memory performance in wild type mice. However, elevated expression of lactate-producing enzymes correlated with poorer memory performance in APP/PS1 mice [36]. In a subsequent study, lactate production was shown to be necessary for acquisition of new spatial memories, but not for retrieval of existing memories [37]. Thus, while the transient production of lactate is beneficial for memory acquisition under normal conditions, higher lactate production, or its inadequate clearance, may be detrimental in AD. Notably, these prior studies did not evaluate whether such age– and disease-related metabolic changes differ between males and females, despite the well-documented sex disparity in AD prevalence and emerging evidence of sex differences in brain glucose metabolism more broadly [7,38].

Lactate has also been implicated in chromatin remodeling. Zhang et al. (2019) identified a new form of epigenetic modification whereby lactate participates in the post-translational alteration of histones with the addition of a lactyl group at lysine residues; a process referred to as histone lactylation [39]. Furthermore, these modifications are sensitive to elevated lactate levels and are linked to regulation of gene expression at distinct locations not activated by histone acetylation 39-41].

Lactate-dependent histone lactylation is specifically increased in microglia adjacent to Aβ plaques in brain tissues from both a 5XFAD mouse model of AD as well as from human post-mortem AD brain samples [42]. Furthermore, lactylated H4 histones at lysine residue 12 (H4K12la) are enriched in promoter regions of glycolytic genes, contributing to their increased expression and related glycolytic activity, resulting in a positive feedback loop driving pro-inflammatory microglial activation along with microglial dysfunction in 5XFAD mice [42]. Chronic neuroinflammation associated with microglial dysfunction is currently believed to be another distinguishing feature of AD [43]. As with lactate metabolism, sex as a biological variable has not been assessed in prior work on microglial histone lactylation, leaving open the question of whether this epigenetic response to amyloid stress differs between males and females. In summary, it has become increasingly evident that lactate is an essential signaling molecule as well as a valuable energy substrate that is part of a flexible but finely tuned metabolic pathway, necessitating further investigation.

Matrix-assisted laser desorption/ionization time-of-flight imaging mass spectrometry (MALDI-TOF IMS) is a powerful, visual, *in situ* technique combining the analytical capabilities of mass spectrometry with traditional histology to reveal the spatial distribution of biomolecules in prepared tissue sections without the need for external labelling [44–47]. The overall procedure involves the coating of tissue sections with an organic compound or matrix, that permits the ionization of individual molecules following exposure to a laser source [45,47–51]. Charged ions are accelerated within an electric field and as they pass through the TOF tube to the detector, the time of flight is measured which allows for the correlated calculation of mass over charge (m/z) for each respective molecule of interest or analyte that is present [51,52]. This information is subsequently sorted and quantified in order to generate a mass spectrum histogram where peak intensity on the y-axis represents the relative abundance of each analyte according to their m/z value along the x-axis [48,52,53]. During measurement, mass spectra are acquired through a series of repetitive laser pulses at predefined x/y coordinates across the entire tissue sample. Thus, the peak intensity for each analyte from each coordinate can be assembled to map out the spatial distribution of each analyte as a heat map [48,53].

Through the use of MALDI-TOF IMS, we analyzed the temporal and spatial changes of brain metabolites, including lactate, glutamate, pyruvate, and citrate, in control and transgenic AD mice at both 6 and 18 months of age, in both male and female animals. In light of the central role of lactate and metabolism in memory formation and AD progression, it was expected that progressive metabolic dysfunction within the hippocampus and cortex of an AD transgenic mouse model would lead to elevated lactate accumulation and altered metabolic flux relative to wild type mice. Additionally, alternate sections from the same brain tissues used for MALDI-TOF IMS analysis underwent immunofluorescent staining using a pan-lysine lactylation antibody to correlate lactate levels with histone lactylation and amyloid deposition in 18-month-old AD mice brains. It was hypothesized that increased histone lactylation levels would co-localize within anatomical brain regions characterized by increased lactate and amyloid accumulation in AD mice, and that this relationship would be modified by sex.

## Materials and Methods

### Mouse model

APPswe/PSEN1dE9, also referred to as APP/PS1 [36,54], was the transgenic mouse model of AD used for this study. These mice express the Swedish mutant form of APP as well as the mutant form (deletion of Exon 9) of presenilin 1 (PSEN1) and were maintained on a C57BL/6 background (Charles River Laboratories International, Saint Constant, QC, Canada). This AD mouse model accumulates amyloid beta (Aβ) by 6 months of age and exhibits cognitive deficits by 9 months [55,56]. Non transgenic litter mates were used as wild type controls. Brain tissues were obtained from both male and female wild type and transgenic mice (n=3 per cohort), at ages 6 and 18 months, for a total of 24 mice.

All mice were housed on a 12 h light/dark cycle with ad libitum access to water and 5015 breeder chow base diet (LabDiet®, Richmond, IN, USA). All procedures were conducted in compliance with the Canadian Council of Animal Care (CCAC) guidelines with approved animal protocols from the Institutional Animal Care and Use Committees at the University of Western Ontario (Protocol number 2020-112).

### MALDI IMS chemicals and sample preparation

The matrix 1,5-diaminonaphthalene (DAN) (Sigma-Aldrich, Oakville, ON, Canada) as a dry weight of 300 mg was applied using the sublimation method [47]. Re-crystallized α-cyano-4-hydroxycinnamic acid (CHCA, Sigma-Aldrich, St. Louis, MO, USA) was used as a calibration standard and was prepared in 50% liquid chromatography grade acetonitrile (ACN, Sigma-Aldrich, St. Louis, MO, USA)/49.9% deionized water (diH_2_0)/0.1% trifluoroacetic acid (TFA, Fisher Scientific, Fairlawn, NJ, USA) at a concentration of 10 mg/ml. The solution was spotted around the tissue on areas with no matrix by applying 0.5 µl aliquots of the solution. Each aliquot was allowed to air dry before a second 0.5 µl aliquot was added directly on top of the initial dried droplet. This CHCA standard served as an external control and its resulting adduct peaks were used for mass calibration prior to each data acquisition session by the mass spectrometer.

Brain tissue was obtained by anesthetizing mice in a CO_2_ chamber followed by euthanasia and transcardial perfusion with phosphate buffered saline (PBS, pH 7.4, Fisher Scientific, Ottawa, ON, Canada) containing protease and phosphatase inhibitors, 1 mM phenyl-methyl-sulfonyl fluoride and 1 mM sodium orthovanadate, respectively (Sigma-Aldrich, St. Louis, MO, USA). The brain was then quickly isolated, cut down the midline along the sagittal plane, and the left hemisphere flash frozen on dry ice. All acquired tissues were subsequently stored at –80 ^0^C until further processing.

Frozen brain samples were mounted to a metal chuck (Fisher Scientific, Ottawa, ON, Canada, catalog number 14-071-414) or specimen holder with diH_2_0 and sectioned on a cryostat at –18 ^0^C (CryoStar NX50, Thermo Fisher Scientific, Burlington, ON, Canada), to a thickness of 10 µm. Tissue sections were thaw-mounted onto conductive glass slides coated with indium tin oxide (ITO) (Hudson Surface Technology, Fort Lee, NY, USA) and stored overnight, at –80 ^0^C, until matrix application the next day.

The application of DAN was based on a prior protocol [47]. Briefly, following removal from overnight storage, the tissue sections mounted on ITO glass slides were dried in a vacuum desiccator for 30 min. The optimal temperature used was 140 ^0^C with a sublimation time of 7 min. Slide-mounted tissue sections were then rehydrated overnight, at –20 ^0^C, and subsequently dried in a vacuum desiccator for another 30 min prior to mass spectrometry analysis.

### MALDI IMS analysis and data processing

All mass spectra and associated images were obtained using an AB Sciex 5800 MALDI TOF/TOF mass spectrometer (Framingham, MA, USA) equipped with a 349 nm Nd:YLF OptiBeam on-axis laser at a pulse rate of 400 Hz and a lateral resolution of 70 µm. Preceding each run, the laser energy was optimized based on the balance between peak resolution and signal-to-noise ratio using the CHCA matrix as a calibration standard at a mass tolerance of +/− 50 ppm. This external calibration, followed by the corresponding data obtained from each respective tissue section, was achieved in negative ion mode. The number of laser shots that were applied for imaging in each pixel was 40. The range of detection was filtered to select for m/z of 70 to 550. Further, data acquisition and processing were accomplished using TOF/TOF Series Explorer and Data Explorer (AB Sciex, Framingham, MA, USA). Final ion distribution data were visualized and interpreted using MSiReader (Version 1.01, FTMS Lab for Human Health Research, North Carolina State Univ, USA). Moreover, through the use of MSiReader, visual images were parsed by selected regions of interest (ROI) and all corresponding ion distribution data, per ROI, were extracted as Excel spreadsheets for further statistical analysis.

### Cryo-immunofluorescent staining and microscopy

Concurrent with brain tissues sectioned for MALDI IMS use, adjacent tissue sections were also collected for dual immunofluorescent (IF) staining. Tissue sections (10 µm thickness) from 18-month-old mice were thaw-mounted onto a Superfrost Plus glass microscope slide (Fisher Scientific, Pittsburg, PA, USA, catalog number 12-550-15).

The affixed tissue sections were then dried on a 45 ^0^C heat block, for 20 min. Subsequently, the tissue sections were post fixed for 1 min, at room temperature, in a solution containing 80% ethanol, 5% glacial acetic acid, and 3.7% formaldehyde (Alfa Aesar, Tewksbury, MA, USA). The samples were rinsed by submerging them in reverse osmosis-treated water followed by 3 x 5 min washes in PBS, pH 7.4 (Electron Microscopy Sciences, Hatfield, PA, USA). A heat retrieval step was next employed in which the slides were placed in pre-heated 10 mM sodium citrate buffer (Sigma-Aldrich, St. Louis, MO, USA), pH 6.0, at 60 ^0^C, and allowed to cool to room temperature (approximately 1.5 h). Once cooled, each tissue section was encircled with a hydrophobic pen (DAKO, Agilent Technologies, Santa Clara, CA, USA).

To quench autofluorescence, samples were placed under UV light for 45 min, then incubated in freshly prepared True-Black (Biotium, Fremont, CA, USA) for 30 sec, at a dilution of 1:20 using 70% ethanol as the solvent. Following a wash with PBS, tissue samples underwent background blocking by incubation in mouse-specific FAB fragments (AffiniPure Fab Fragments Goat Anti-Mouse IgG, Jackson Immuno Research, West Grove, PA, USA, product code 115-007-003) diluted to 1:35 in PBS, for 60 min, followed by an application of Background Sniper (Biocare Medical, Pacheco, CA, USA) for 7 min. Two primary antibodies were freshly prepared together in DAKO Universal Antibody Diluting solution (Agilent Technologies, Mississauga, ON, Canada). A rabbit anti-L-lactyllysine antibody (PTM Biolabs, Chicago, IL, USA, catalog number PTM-1401) was used at a dilution of 1:200 while a mouse anti-beta amyloid antibody (Novus Biologicals, Toronto, ON, Canada, catalog number NBP2-13075) was used at a dilution of 1:500. The brain sections were incubated overnight, at 4 ^0^C, within a humidity chamber.

The next day, slides underwent 3 x 5 min washes with PBS followed by incubation with goat, anti-rabbit Alexa Fluor 647 and goat, anti-mouse Alexa Fluor 568 (Thermo Fisher Scientific, Mississauga, ON, Canada) secondary antibodies at a dilution of 1:500 in DAKO, for 40 min, at room temperature. Brain sections were then counterstained with DAPI (Thermo Fisher Scientific, Mississauga, ON, Canada, catalog number D1306) for 2 min, at a dilution of 1:300 in PBS. After adding coverslips, the affixed tissue sections were sealed with anti-fade Prolong-Gold mounting medium (Thermo Fisher Scientific, Mississauga, ON, Canada) and the sealed slides were left to cure for 24 h, in the dark, at room temperature. The sealed, tissue mounted slides were then stored at 4 ^0^C, in the dark, until time of imaging.

Full scans of each sagittal brain section were obtained at 20x magnification on a Nikon inverted Ti2E deconvolution microscope (Nikon Canada Inc., Mississauga, ON, Canada) using the large image stitching method (Nikon’s NIS Elements, Mississauga, ON, Canada). Two distinct regions, highlighting the hippocampus and cortex, were then selected, cropped from each full brain image, saved as ND2 files, and exported as TIFF files for further statistical analysis. Monochrome images of each separate channel along with open microscopy environment (OME) metadata were included with exported files.

### Statistical analysis

GraphPad Prism (version 9.4.1) was used to generate all graphs and perform statistical analyses with the exception of the principal component analysis (PCA) and its associated graphs. Based on age, sex, and genotype, a three-way analysis of variance (ANOVA) followed by Tukey’s post hoc test was conducted to evaluate changes in metabolite levels and bi-directional ratios related to metabolic flux. In addition, a two-way ANOVA was utilized for assessing lactylation activity between genotype and sex within each of three intensity categories as well as lactylation activity based on distance to plaques and sex in transgenic mice only. Parametric analysis was done and the data was observed to be non-parametric. Thus, Spearman correlation analysis was next performed to compare the association between lactate level and corresponding lactylation activity found within the same anatomical brain regions. XLSTAT Premium version (Addinsoft, Paris, France) was used for PCA to ordinate the changes in anatomical regions that co-segregated with the altered metabolite levels, metabolic flux, and lactylation activity. Visualization of the biplot as well as the associated univariate plots were also performed with this software.

In order to first normalize the ion distribution data obtained through MALDI IMS, area under the curve (AUC) for each individual metabolite of interest per ROI was obtained using Excel and this value was further taken as a ratio to the AUC of the mass spectrum for the entire ROI in which each respective metabolite was found. The full mass spectrum AUC for each ROI was determined using GraphPad.

For data extraction from TIFF files of immunofluorescence images into an Excel spreadsheet, Image-Pro (version 10) was programmed to capture percent area and integrated optical density (IOD), at three different levels of signal intensity, when comparing entire ROIs between APP/PS1 transgenic and control brain sections. Raw IOD data were further normalized by taking the sum of all IOD present in a selected ROI and dividing by the sum of the area in pixels to obtain the mean gray value. Additionally, for analysis of proximal versus distal regions surrounding plaques in transgenic brain tissue, Image-Pro (version 11) was programmed to extract the data directly from the respective ND2 files.

Furthermore, amyloid plaques were scored by utilizing both an intensity and size filter. Amyloid-based signal was determined by employing *Otsu’s minimum variance threshold method* and any signal greater than 50 pixels in size was automatically identified as a plaque. These parameters worked well in the cortex and *dentate gyrus* ROIs where amyloid burden was greatest, but some tissue samples required a manual adjustment to the upper limit of amyloid signal detected in order to score some plaques for sampling in the CA1 and CA2/CA3 subregions. Once plaque(s) were detected, a surrounding region was demarcated that spanned out by 32 pixels, or by approximately a 10 µm radius, from the edge of a plaque and was defined as the “proximal” region while anything outside the proximal region, but still within 100 pixels, or a 32 µm radius from the edge of a plaque, was considered the “distal” region.

## Results

### MALDI-TOF IMS assessment of brain metabolite content using a DAN matrix Spatial and temporal distribution of individual metabolites

MALDI IMS analysis of individual metabolites was performed using DAN as the matrix. The normalized AUC was calculated for each peak specific to lactate, glutamate, pyruvate, and citrate, acquired from 9 different brain regions **(Figure S1)**: **(1)** the CA1 subregion of the hippocampus, **(2)** the combined CA2 and CA3 subregions of the hippocampus, **(3)** the *dentate gyrus* (DG) hippocampal subregion, **(4)** the cortex, **(5)** the thalamus, **(6)** the cerebellum, **(7)** the white matter of the cerebellum, **(8)** the corpus callosum, and the entire brain section. Initially, a three-way ANOVA followed by Tukey’s post hoc test was conducted for each metabolite, per ROI, based on age, sex, and genotype. Most noteworthy changes in metabolites occurred in the three subregions of the hippocampus as well as in the thalamus, thus further analysis focused on these regions.

Levels of glutamate were highest in all sampled ROIs; at approximately three times the amount of lactate **(Figures 1A, 2A)**. In comparison to lactate, both pyruvate and citrate levels were considerably lower by a factor of ten **(Figures 3A, 4A)**.

**Figure 1:**
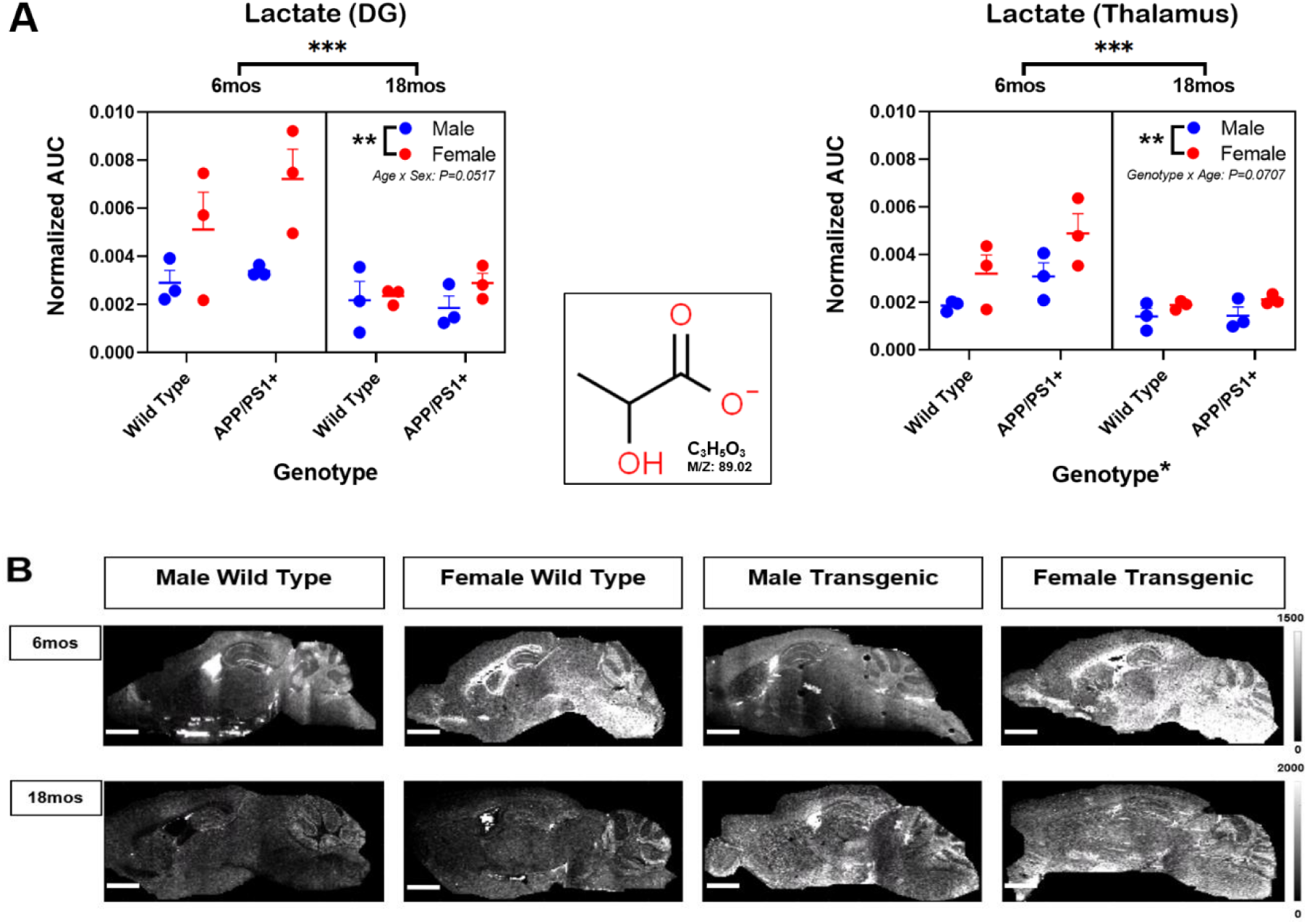
Regional levels of lactate in young versus aged mouse brains. Lactate levels, as determined by MALDI IMS, revealed a significant decline with age for both wild type and APP/PS1 mice for both sexes (**A**). Lactate levels were significantly higher in female mice compared to their male counterparts for both age groups. A genotype-based effect was confirmed in the thalamus whereby APP/PS1 transgenic mice had higher levels of lactate than their respective wild type litter mates which was more apparent in the 6-month-old age group. Heat maps revealed that although lactate was prevalent throughout many ROIs, the signal was stronger in white matter regions, at 6 months of age, for both female cohorts but decreased with age (**B**). *\* P < 0.05, ** P < 0.01, *** P < 0.001*. Individual values obtained for each mouse are graphed with mean ± SEM (n=3); scale bars on IMS images represent 20 mm; lateral resolution of 70 µm [Molecular formula and structure obtained from ^©^ChemSpider, courtesy of Royal Society of Chemistry, UK].

Changes in metabolite levels as a function of age, sex, and genotype were also assessed. As expected, glutamate and lactate levels varied in parallel with one another **(Figures 1A, 2A)**. Overall metabolite levels were higher, for both sexes, at 6 months of age, and significantly decreased at 18 months of age. Furthermore, females tended to have significantly elevated levels of each metabolite in comparison to their male counterparts. Within each age group, the interaction of both age and sex was confirmed to have some influence on lactate levels in the DG (F=4.4190, P=0.0517) along with having significant effects on glutamate levels in the thalamus (F=5.3480, P=0.0344). Interestingly, the ROI where genotype had a significant impact was in the thalamus, with elevated lactate levels observed in transgenic mice compared to wild type mice at 6 months of age (F=5.3030, P=0.0350).

Of note was the degree of variability within each cohort. With respect to lactate, there was a much higher degree of variability for the 6-month-old females as opposed to the 6-month-old male groups **(Figure 1A)**. However, the opposite was true for glutamate levels whereby there was a higher degree of variability amongst 6-month-old males compared to females at the same age **(Figure 2A)**. The degree of variability noticeably decreased with age and the levels were much more consistent at 18 months of age for both metabolites **(Figures 1A, 2A)**.

**Figure 2:**
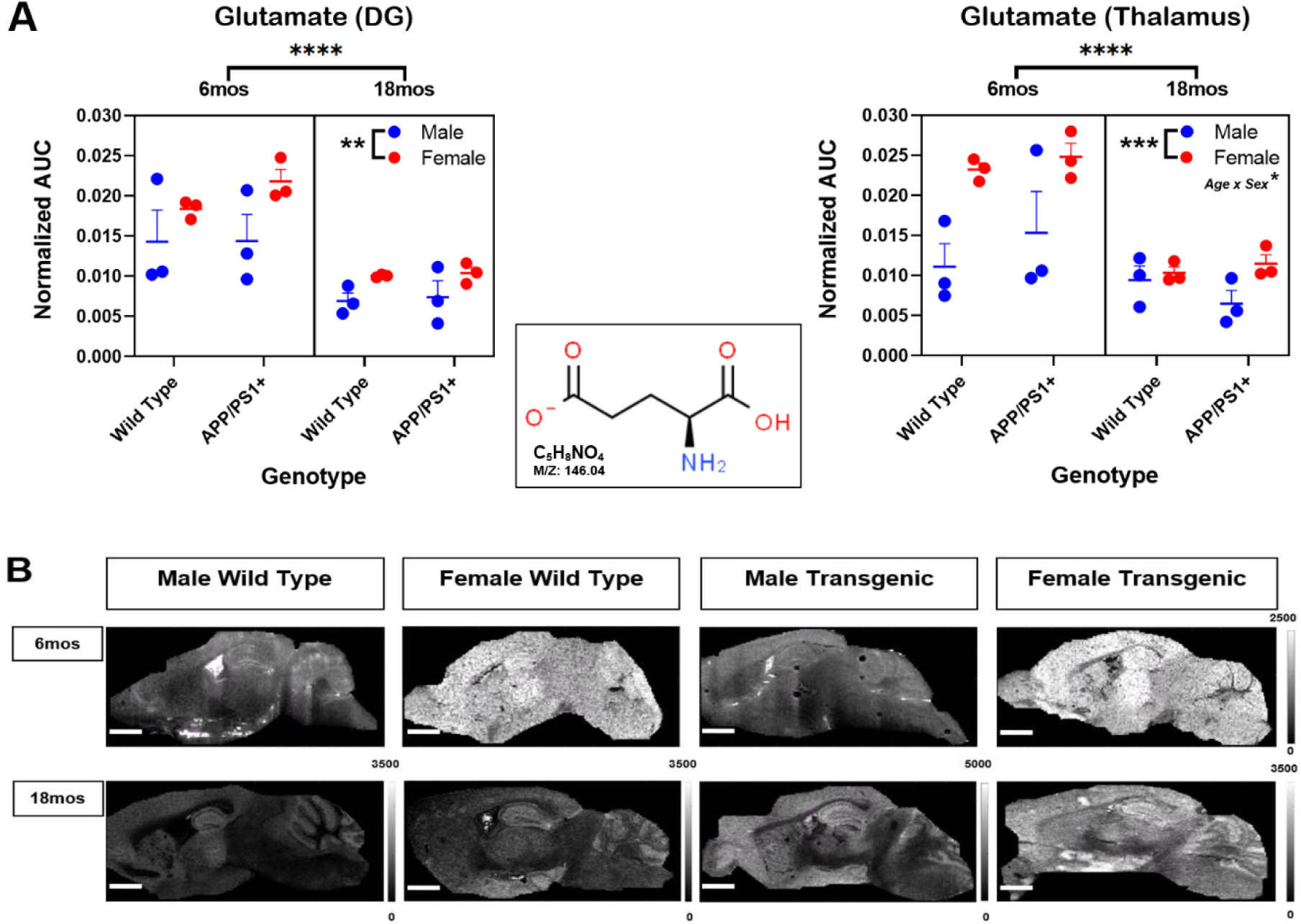
Regional levels of glutamate in young versus aged mouse brains. Glutamate levels assessed by MALDI IMS were elevated in all sampled regions, at approximately three times the amount of lactate (**A**). Glutamate levels were significantly higher in young females compared to young males but declined with age for both sexes. The interaction of age and sex had a significant effect in the thalamus. Glutamate was the dominant signal in brain tissue at both age groups and was readily detected in grey matter regions such as the cortex, hippocampus, thalamus, and cerebellum (**B**). *\* P < 0.05, ** P < 0.01, *** P < 0.001, **** P < 0.0001*. Individual values obtained for each mouse are graphed along with mean ± SEM (n=3); scale bars on IMS images represent 20 mm; lateral resolution of 70 µm [Molecular formula and structure from ^©^ChemSpider, courtesy of Royal Society of Chemistry, UK].

**Figure 3:**
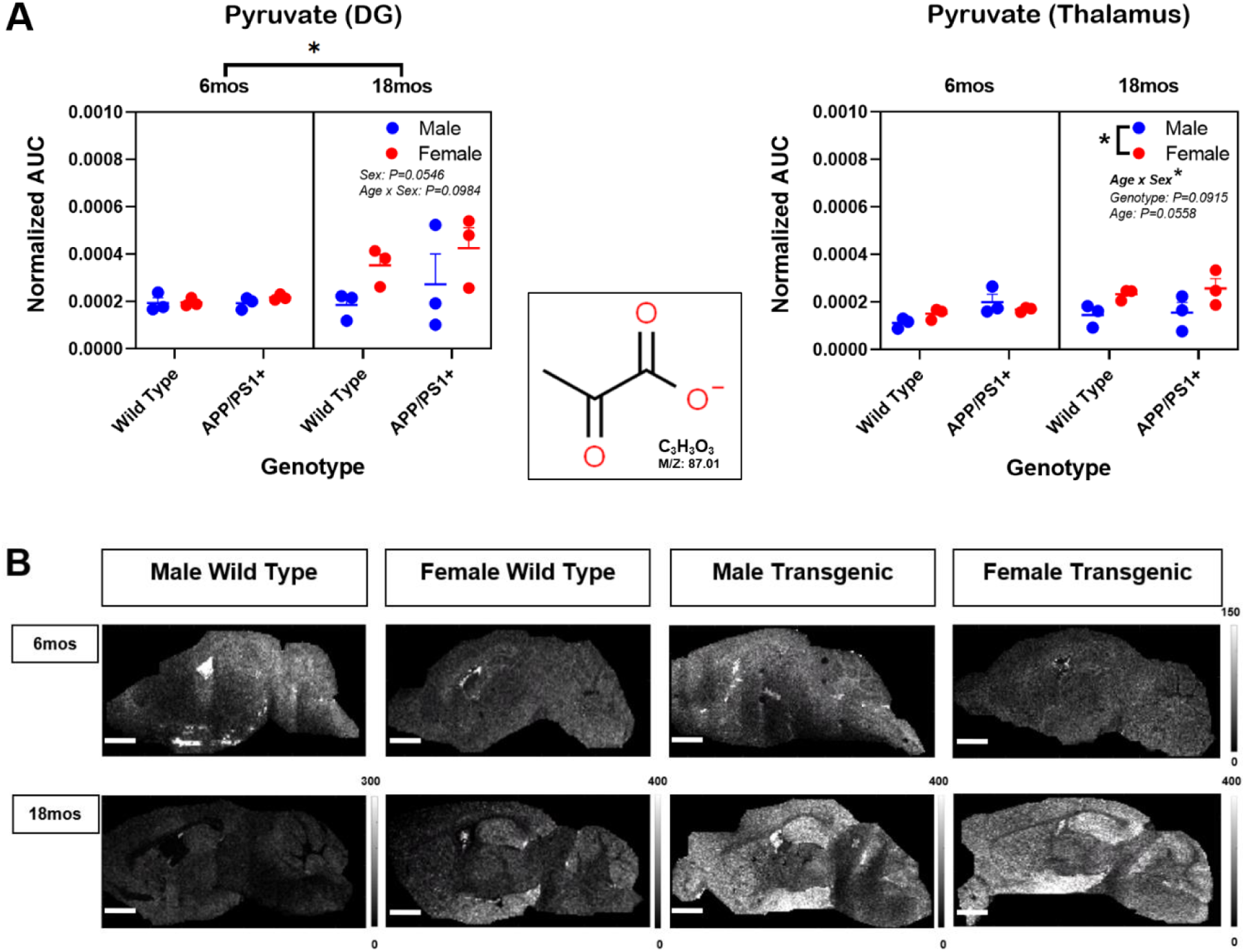
Regional levels of pyruvate in young versus aged mouse brains. MALDI IMS analysis revealed that pyruvate levels were considerably lower than lactate levels in all brain regions of interest (**A**). Pyruvate levels increased in the brains of aged female mice compared to young female mice, regardless of genotype. This increase was reflected in IMS images revealing elevated pyruvate within the hippocampus and cerebellum (**B**). *\* P < 0.05.* Individual values obtained for each mouse are graphed along with the mean ± SEM (n=3); scale bars on IMS images represent 20 mm; lateral resolution of 70 µm [Molecular formula and structure obtained from ^©^ChemSpider, courtesy of Royal Society of Chemistry, UK].

With respect to variability amongst cohorts, pyruvate levels were much more consistent, at 6 months of age, for all male and female groups **(Figure 3A)**. However, the degree of variability increased and was much more pronounced for both the male and female transgenic mice, at 18 months of age, in the DG. In addition, there was an age effect whereby pyruvate levels were significantly higher at 18 months of age compared to younger mice. Further, pyruvate levels in the DG showed an effect of sex (F=4.3000, P=0.0546), while both sex (F=6.3800, P=0.0225) and the interaction of sex with age (F=5.4210, P=0.0333) were found to have significant effects in the thalamus.

Citrate levels exhibited variability amongst all groups, particularly at 6 months of age **(Figure 4A)**. Mean citrate levels were higher in males compared to females, in younger mice, but this trend reversed at 18 months of age whereby higher citrate levels were observed in females. Additionally, within each age group, mean citrate levels were higher in wild type females compared to transgenic females and higher citrate levels were observed in transgenic males versus wild type males. However, many of these differences were not deemed to be significant.

**Figure 4:**
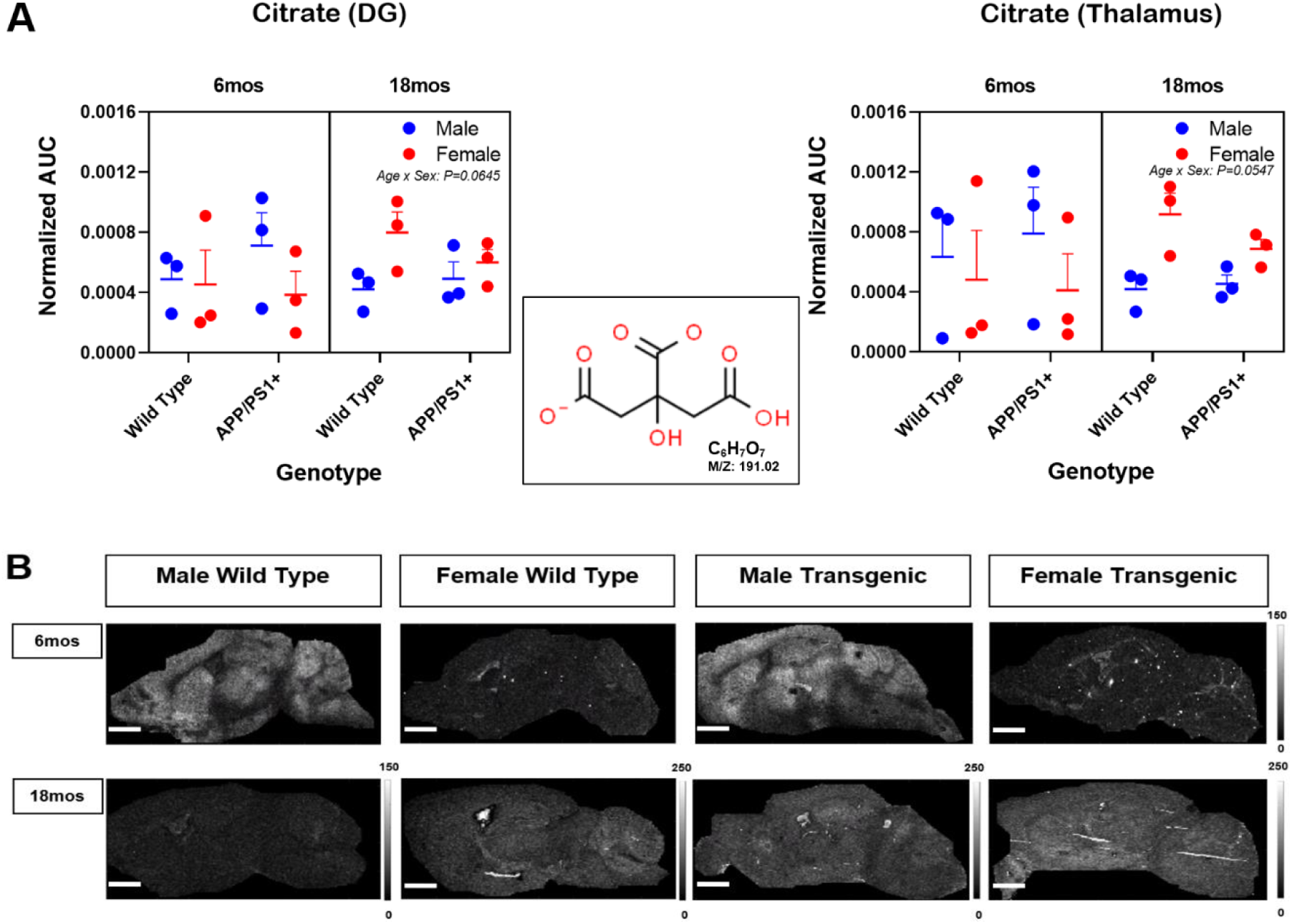
Regional levels of citrate in young versus aged mouse brains. MALDI IMS was employed to detect citrate levels in the hippocampal and thalamic regions of the brain (**A**). Factors of age, sex, and genotype were found to have no significant influence on citrate levels in sampled ROIs. IMS images further highlighted these levels tended to be higher in young males but increased in aged females (**B**). *\* P < 0.05*. Individual values obtained for each mouse are graphed along with the mean ± SEM (n=3); scale bars on IMS images represent 20 mm; lateral resolution of 70 µm [Molecular formula and structure obtained from ^©^ChemSpider, courtesy of Royal Society of Chemistry, UK].

MALDI-TOF IMS data was also processed to generate heat maps highlighting the spatial and temporal levels of individual metabolites **(Figures 1B, 2B, 3B, 4B)**. A representative set of female and male brain tissue, at 6 and 18 months of age, revealed the regional distribution and relative intensity levels for each metabolite including lactate, glutamate, pyruvate, and citrate. Glutamate was the most abundant in 6-month-old females and was readily detected in grey matter regions such as the cortex, hippocampus, thalamus, and cerebellum **(Figure 2B)**. Glutamate continued to be the dominant signal in the 18-month-old transgenic female. Interestingly, while lactate was prevalent throughout many ROIs, the signal was stronger in white matter regions, at 6 months of age, for both female transgenic APP/PS1 and wild type mice as opposed to their male counterparts **(Figure 1B)**. However, lactate levels in the white matter regions decreased with age in the 18-month-old mice. While 6-month-old male mice appeared to have slightly elevated pyruvate and citrate levels in comparison to 6-month-old female mice, there was a discernable increase in both pyruvate and citrate levels for both 18-month-old transgenic and wild type female mice **(Figures 3B, 4B)**.

### Spatial and temporal assessment of metabolic flux

The direction of metabolic flux was assessed by comparing ratios of pyruvate to lactate **(Figure 5A & B)**, representing changes in glycolysis and lactate production, versus ratios of pyruvate to citrate **(Figure 5C & D)**, reflective of entry into the TCA cycle and changes in OXPHOS. Both directional-based ratios were assessed for any significance by employing a three-way ANOVA, per ROI, based on age, sex, and genotype.

**Figure 5:**
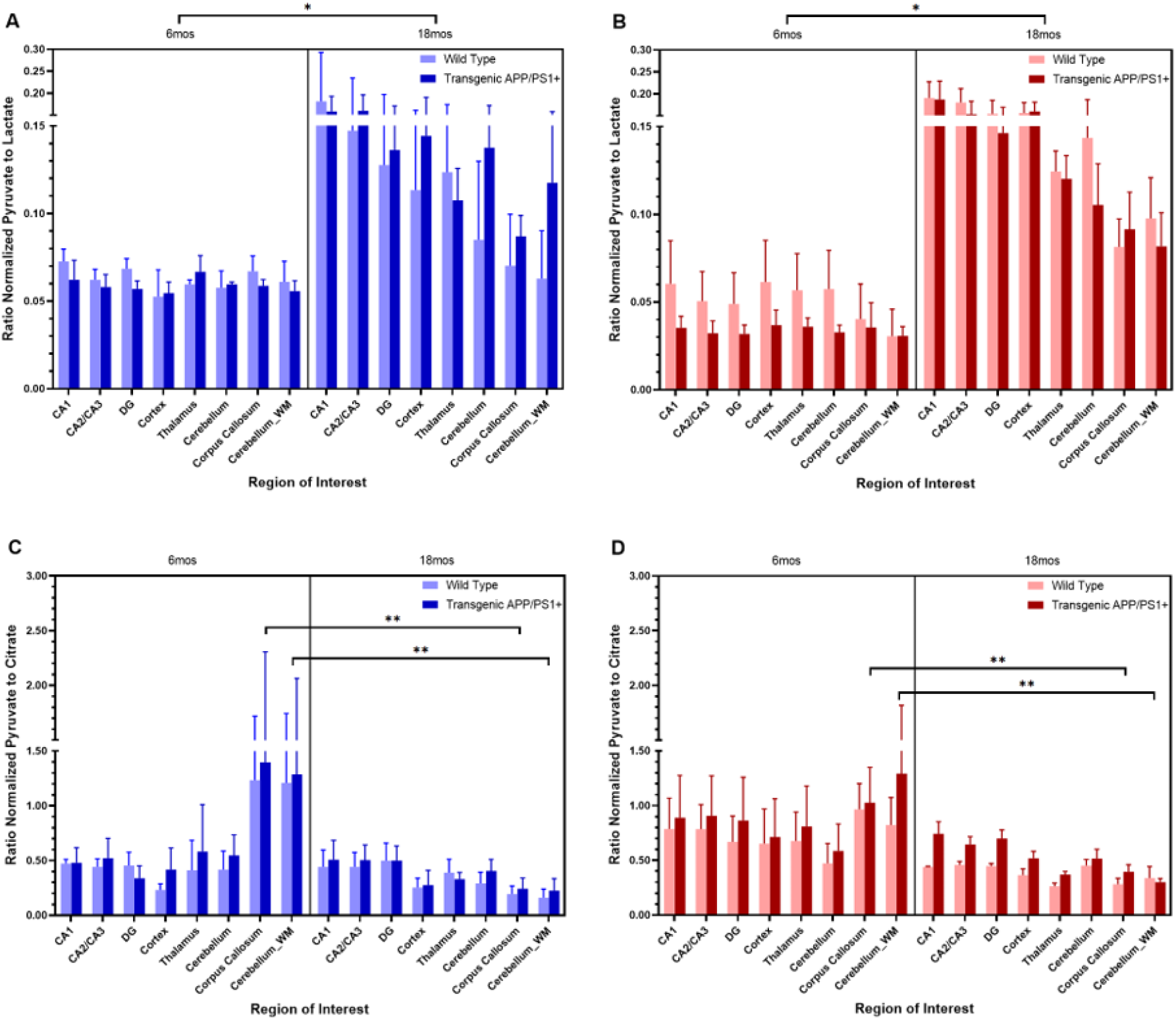
Alterations of metabolic flux in young and old wild type and transgenic AD mice. The direction of metabolic activity was assessed by determining the ratio of pyruvate to lactate, representing changes in glycolysis and lactate production, versus the ratio of pyruvate to citrate, characterizing entry into the TCA cycle and changes in OXPHOS. Pyruvate to lactate ratios significantly increased with age in both male (**A**) and female (**B**) cohorts, indicating a shift away from lactate production in 18-month-old mice. In contrast, pyruvate to citrate ratios in both male (**C**) and female (**D**) cohorts revealed a significant reduction in white matter regions with age. *\* P < 0.05, ** P < 0.01*. Each bar represents the mean value ± SEM (n=3).

There was a significant two-to three-fold increase in pyruvate to lactate ratios for both male **(Figure 5A)** and female **(Figure 5B)** 18-month-old mice, in comparison to 6-month-old mice across all 8 brain regions; indicating a shift away from glycolysis and less lactate being produced or consumed. Ratios reflected an approximate 1:20 pyruvate to lactate relationship in 6-month-old mice as opposed to a 1:8 ratio in the grey matter regions of 18-month-old males or 1:5 for the older females within the same regions. However, there were no significant differences found based on genotype, sex, or any combination of the three independent factors. Nonetheless, mean pyruvate to lactate ratios tended to be lower in the grey matter regions of the younger female, transgenic mice, indicating more glycolytic activity than the female wild type group.

In contrast, there was a significant decrease in pyruvate to citrate ratios in the white matter regions of both aged male **(Figure 5C)** and female mice **(Figure 5D)** (Corpus Callosum: F=9.9080, P=0.0062; Cerebellum WM: F=10.2000, P=0.0056). The pyruvate to citrate relationship approached a 3:2 ratio in the white matter regions of the younger mice but dropped to a 1:2 ratio in the older mice.

Co-localized MALDI IMS images of lactate and citrate were assessed to gain a visual perspective of the metabolic flux in the brain; regions highlighting the presence of lactate representing glycolytic activity in red contrasted with regions of pronounced citrate presence representing regions of movement in the TCA cycle and associated OXPHOS activity in blue **(Figure 6)**. In young female mice, the elevated presence of lactate was observed throughout the brain, but glycolytic activity was more pronounced in female transgenic compared to wild type mice for most regions, such as the cortex, hippocampus, thalamus, and cerebellum. Additionally, co-localization with citrate was more apparent in the neuron dense DG region of the hippocampus and cerebellum of the 6-month-old female transgenic mouse as visualized by a magenta colour.

**Figure 6:**
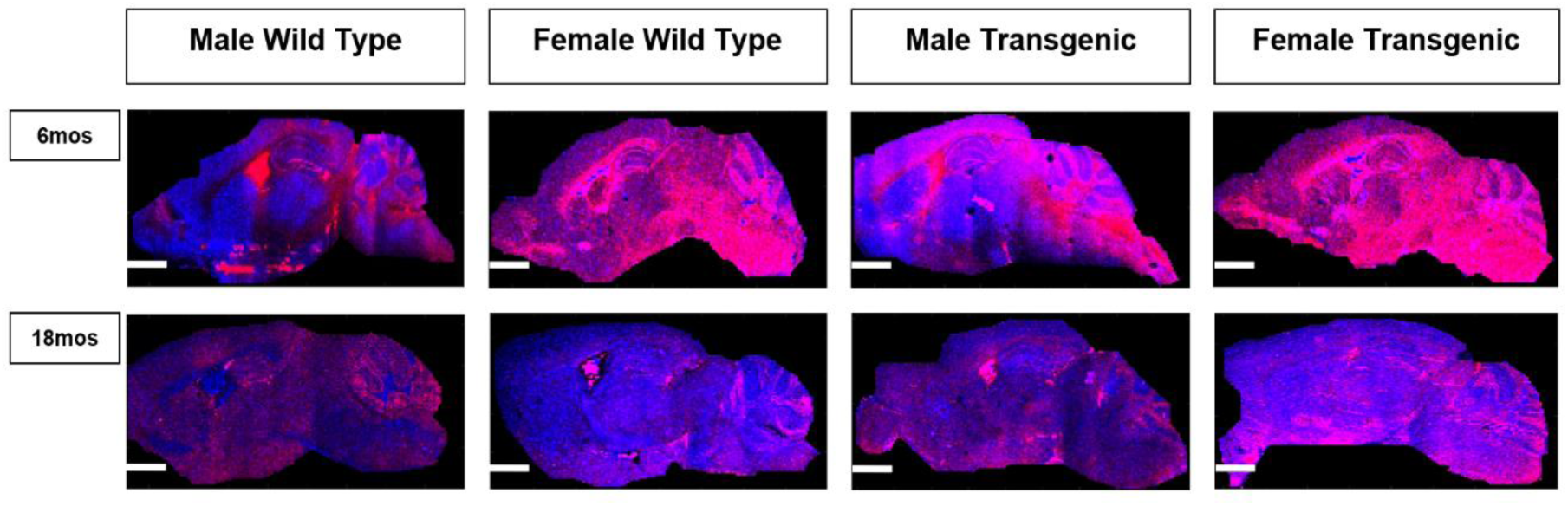
Co-localized MALDI IMS images of lactate and citrate highlighting regions of glycolytic activity compared to regions of oxidative phosphorylation. A visual perspective of metabolic flux throughout the brain was obtained by combining IMS images of relative lactate and citrate distribution. The presence of lactate reflecting glycolytic activity is shown in red while the presence of citrate representing movement in the TCA cycle and associated OXPHOS is in blue. Any co-localization of the two metabolites appears as magenta in colour. Widespread glycolytic activity was present in both samples of young female mice, but as lactate levels decreased with age, OXPHOS activity became more pronounced in many regions. The reverse was true for males and a lower level of glycolytic activity was maintained at 18 months of age with the exception of the hippocampus in the 18-month-old transgenic male where prominent OXPHOS activity was present. Scale bars on IMS images represent 20 mm; lateral resolution of 70 µm.

Within the brain tissue of the 6-month-old male mice, citrate levels were much more pronounced and OXPHOS activity was detected in the cortex, hippocampus, thalamus, and cerebellum. However, there was much variation within these regions as well. For example, there appeared to be more OXPHOS activity in the less neuron dense region of the DG, also referred to as the molecular layer, compared to the less neuron dense area in the CA1. Further, there was increased co-localization occurring in the cortex and neuron dense, granular layers of both the DG and cerebellum in the 6-month-old transgenic mouse. As levels of lactate production or consumption decreased with age and citrate levels increased in both female cohorts, OXPHOS activity appeared to be more prominent in the hippocampus, thalamus, and cerebellum of the 18-month-old female wild type mouse compared to the female APP/PS1 transgenic mouse. Moreover, in both 18-month-old male cohorts, a lower level of glycolytic activity was observed throughout the brain, but there was some variation in activity across the cortex and elevated OXPHOS activity was observed in the hippocampus of the transgenic male mice.

### Principal component analysis of metabolites and metabolic flux in young versus aged mice

MALDI IMS-based data obtained from all 24 mouse brain tissue samples was re-examined by way of principal component analysis (PCA). Differences in cerebral metabolite levels and metabolic flux were determined based on all four factors of genotype, age, sex, and brain region **(Figure S2A-D)** with the output presented in the biplot accounting for 70.03% of the total variability in the data. Levels of lactate, glutamate, pyruvate, and the directional ratio of pyruvate to citrate were predominantly associated with the younger 6-month-old mice within anatomical brain regions that included the CA2/CA3 and DG subregions of the hippocampus along with the full brain **(Figure S2A-D, top right quadrants)**. Citrate and the directional ratio of pyruvate to lactate were predominantly associated with the aged 18-month-old mice within the CA1 subregion of the hippocampus **(Figure S2A-D, top left quadrants)**. There was no clear pattern of association present based on genotype.

Two sample t-tests were applied to determine the distinct impact for each of genotype, age, and sex **(Figure 7A-C)**. Levels of both lactate (p=0.035) and pyruvate (p=0.010) were significantly higher in transgenic mice compared to wild type mice **(Figure 7A)**. In addition, lactate (p < 0.0001) and glutamate (p < 0.0001) levels were significantly elevated in 6-month-old mice compared to 18-month-old mice **(Figure 7B)**. In contrast, citrate levels were significantly elevated in the aged mice compared to the younger mice (p=0.009). Age also had a notable effect on both directional ratios whereby pyruvate to lactate levels were significantly elevated (p < 0.0001) in 18-month-old mice compared to the younger cohort; and pyruvate to citrate levels were elevated in the 6-month-old mice compared to their aged counterparts. Additionally, increased metabolite levels were evident in females compared to males. Specifically, lactate (p < 0.0001), glutamate (p < 0.0001), pyruvate (p=0.001), and pyruvate to citrate levels (p=0.019) were significantly higher in female mice compared to male mice **(Figure 7C)**.

**Figure 7:**
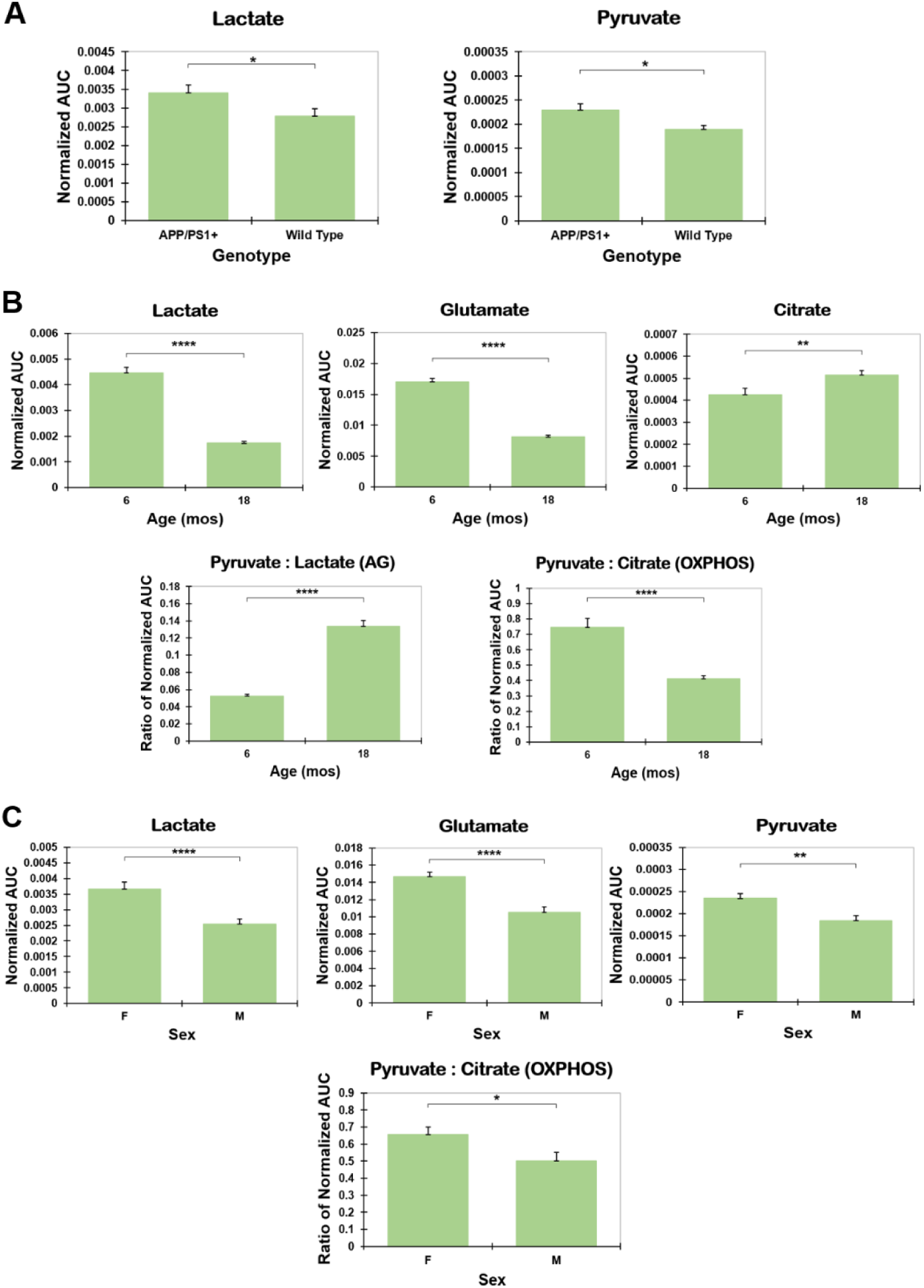
Two sample t-tests based on the discrete factors of genotype, age, and sex. A simplified analysis based on each independent variable revealed that lactate was significantly highest in transgenic mice (**A**), at 6 months of age (**B**), and more so in females (**C**). Additionally, genotype and sex had a significant effect on pyruvate levels whereby levels in transgenic mice (**A**) and specifically females (**C**) were higher compared to wild type and male mice, respectively. In parallel to lactate levels, glutamate levels were significantly elevated in the younger mice (**B**) and in females compared to males (**C**). Alternately, citrate levels were significantly elevated in the aged mice (**B**). Age significantly impacted both directional ratios highlighting higher lactate production or consumption in the young mice compared to the aged mice while the aged mice exhibited increased entry into the TCA cycle and OXPHOS activity. *\* P < 0.05, ** P < 0.01, **** P < 0.0001*. Each bar represents the mean value ± SEM (n=108).

### Cryo-immunofluorescent staining for lactylation

#### Analysis of histone lactylation based on genotype and sex

Lactate accumulation within cells promotes a post-translational modification to proteins, including nuclear histones, termed lactylation [39–42]. In light of the age– and sex-specific changes in cerebral lactate levels observed with both three-way ANOVA and PCA, it was of interest to examine protein lactylation levels in the three subregions of the hippocampus, **(1)** CA1, **(2)** CA2/CA3, and **(3)** DG, as well as in the **(4)** cortex in both 18-month-old wild type and transgenic mice **(Figure S1)**. Brain tissues were stained with an antibody that specifically recognizes lactyl-groups on lysine residues (pan-Kla) followed by analysis using deconvolution fluorescence microscopy. Due to variation in lactylation fluorescence intensity of stained brain tissues, quantitative analysis was divided into “Bright” signal corresponding to an intensity threshold of greater than 6000 and “Dim” signal limited to a range in intensity between 2500 to 5999. Comparison of lactylation staining versus nuclear staining with DAPI revealed that the “Bright” lactylation signal was predominantly localized to small, nucleated cells typical of microglia whereas the “Dim” lactylation signal was found in larger nuclei typically found in neurons **(Figure S3)**. Although additional cell-type specific antibodies were not utilized at this time, delineating staining intensity may provide some insight into cell types exhibiting differential histone lactylation.

There were no significant genotype-based differences in lactylation levels found within any of the four ROIs, at all three intensity levels **(Data not shown)**. However, a sex-based effect was identified within the cortex and CA2/CA3 subregion whereby the proportion of lactylation signal detected in the selected ROIs encompassed a significantly larger area in aged females than in aged males. Further, the brain tissues from female mice exhibited a significantly higher mean gray value or average intensity level of lactylation signal compared to male mice.

### Analysis of histone lactylation in transgenic mice based on distance to amyloid plaques

To determine whether histone lactylation was increased in regions adjacent to amyloid plaques, immunofluorescence intensity was quantified in dual-stained brain tissue. Lactylation signal was assessed within a proximal “neighborhood” surrounding amyloid plaque(s) and compared with signal in regions distal to the same plaque(s), using both percent area positive for lactylation and mean gray value as measures of intensity. Data were analyzed for total lactylation signal and further categorized as “Bright” or “Dim” signal to reflect putative non-neuronal and neuronal cell populations, respectively.

Examining factors of sex and distance to plaque(s) at each intensity level, there was both a sex– and distance-based effect identified in the cortex for all lactylation signal (Sex, F=10.9400, P=0.0107; Distance, F=22.4200, P=0.0015) and the putative non-neuronal, “Bright Only” level (Sex, F=9.8220, P=0.0139; Distance, F=10.4600, P=0.0120) along with the putative neuronal, “Dim Only” intensity level (Sex, F=9.1760, P=0.0163; Distance, F=20.0800, P=0.0021), whereby the percent area was significantly higher in females compared to males and significantly more widespread in proximal regions compared to distal regions for both sexes **(Figure 8A)**. Interestingly, a trend of higher percent area of lactylation proximal to plaques was observed in the female brain tissues for all ROIs and at all intensity levels with the exception of the “Bright Only” signal in the DG.

**Figure 8:**
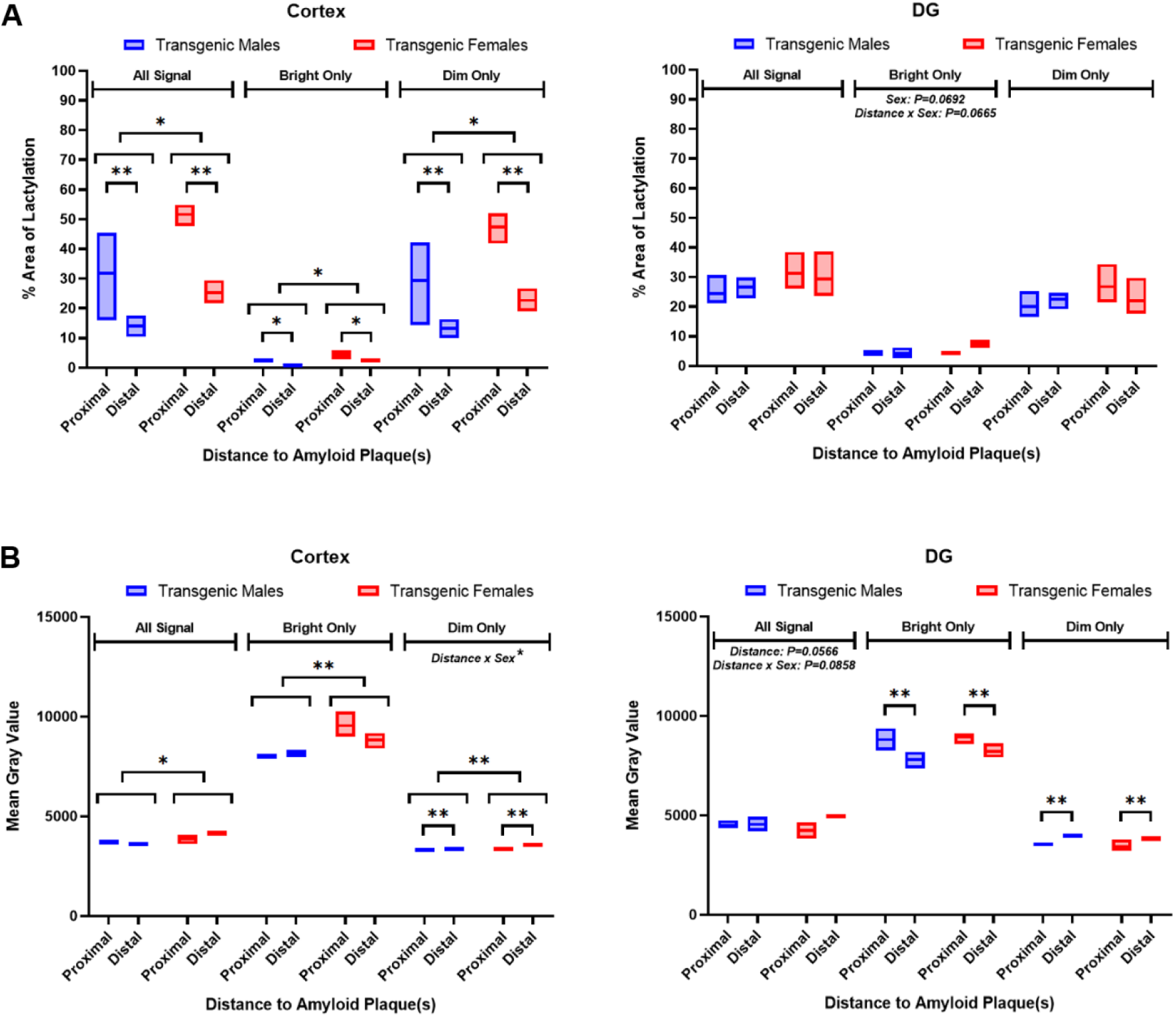
Histone lactylation based on proximity to amyloid plaque(s) in both 18-month-old transgenic male and female mice. Based on two-way ANOVA examining factors of sex and distance to plaque(s) in transgenic mice, the percent area of lactylation was significantly higher in females compared to males at all intensity levels in the cortex, but lactylation was significantly more pronounced in proximal regions rather than distal regions for both sexes (**A**). The degree of variability decreased when mean gray values were being assessed (**B**). Females had significantly higher mean gray values than males, at all intensity levels, in the cortex. The cortex also revealed distance-based and interaction effects in the “Dim Only” category. Average intensity levels of lactylation signal were significantly higher in distal regions of the cortex and DG at the “Dim Only” level, for both sexes, whereas the reverse effect was observed for the “Bright Only” level, in the DG, for both sexes. *\* P < 0.05, ** P < 0.01*. Each floating bar represents the range between the minimum and maximum calculated values obtained for each cohort with a solid horizontal line indicating the mean (n=3).

When examining mean gray values across ROIs, the degree of variability within groups appeared to decrease and values were much more consistent at all intensity levels **(Figure 8B)**. Sex was a significant contributing factor at all intensity levels in the cortex (All Signal, F=10.5200, P=0.0118; Bright Only, F=22.2800, P=0.0015; Dim Only, F=20.0900, P=0.0021). In addition, a distance-based effect was confirmed at the “Dim Only” level in the cortex (F=19.8300, P=0.0021) along with an interaction effect (F=7.7040, P=0.0241) whereby both sexes had significantly higher mean gray values distal to plaques, but the lactylation signal in female tissues was significantly brighter or at an increased level of intensity.

Mean gray values were significantly higher distal to plaques as had occurred in the putative, neuronal “Dim Only” category within the cortex (F=16.1800, P=0.0038). Alternately, lactylation signal was significantly more intense proximal to plaques at the putative, non-neuronal “Bright Only” level (F=12.7800, P=0.0072). Interestingly, this reverse trend of increased lactylation levels observed in proximal regions to plaques within putative non-neuronal cells in the DG, was also consistently present for females within the other three ROIs.

To examine the spatial relationship between amyloid plaques and histone lactylation, brain tissues from both 18-month-old male and female transgenic mice were dual stained with antibodies specific for amyloid and protein lactylation **(Figure S4)**. The amyloid burden was greater in the cortex **(Figure S4A)** and in the DG subregion of the hippocampus **(Figure S4B)**. Additionally, when comparing the individual channels for amyloid and lactylation signal in the transgenic female tissue, lactylation signal was slightly higher and/or more condensed where plaques were present. Overall, the lactylation signal was also slightly higher in the female transgenic tissue.

### Principal component analysis of protein lactylation in aged mice

PCA was used to determine the ordination of protein lactylation in brain tissues of aged mice **(Figure S5)**. Genotype and sex were considered in addition to lactylation across 4 anatomical brain regions. Percent area of lactylation was observed to be ordinated with the hippocampal DG subregion in wild type females **(Figure S5A, top right quadrant)** while the mean gray value ordinated with the bright lactylation signal **(Figure S5A, top left quadrant)**. Furthermore, lactylation signal was also investigated specifically in the brain tissues of aged transgenic mice based on sex and distance to amyloid plaques. The percent area of lactylation was associated with all lactylation signal proximal to observed plaques within the cortex and CA2/CA3 hippocampal subregion **(Figure S5B, top right quadrant)**. However, the mean gray value was specifically associated with the bright lactylation signal found within the DG of the female mice **(Figure S5B, top left quadrant)**.

### Comparison of lactate levels assessed by MALDI-TOF IMS to lactylation detected by IF

A correlational analysis was conducted to compare the lactate levels calculated from MALDI-TOF IMS data to lactylation levels quantified through immunofluorescence microscopy. The calculated values obtained from 18-month-old male and female mouse brain tissues were combined to increase the sample size to 6 per genotype. Lactate levels were compared to the percent area of lactylation as well as to mean gray values obtained within the cortex, CA1, CA2/CA3, and DG, at two intensity ranges: the putative non-neuronal, “Bright” level and the putative neuronal, “Dim” level.

When comparing lactate levels to the percent area of lactylation, there were no significant associations present for both genotypes within each respective ROI **(Data not shown)**. However, when lactate levels were compared to the associated mean gray values within the same ROI **(Figure 9)**, a positive correlation was observed in the cortex (Spearman r = 0.8857, P=0.0333), the CA2/CA3 subregion (Spearman r = 0.8286, P=0.0583), and in the DG (Spearman r = 0.7714, P=0.1028), at the putative non-neuronal, “Bright” intensity level in wild type mice. In transgenic mice, the positive association between the two variables was not apparent in the CA2/CA3 and DG subregions or was weaker in the cortex.

**Figure 9:**
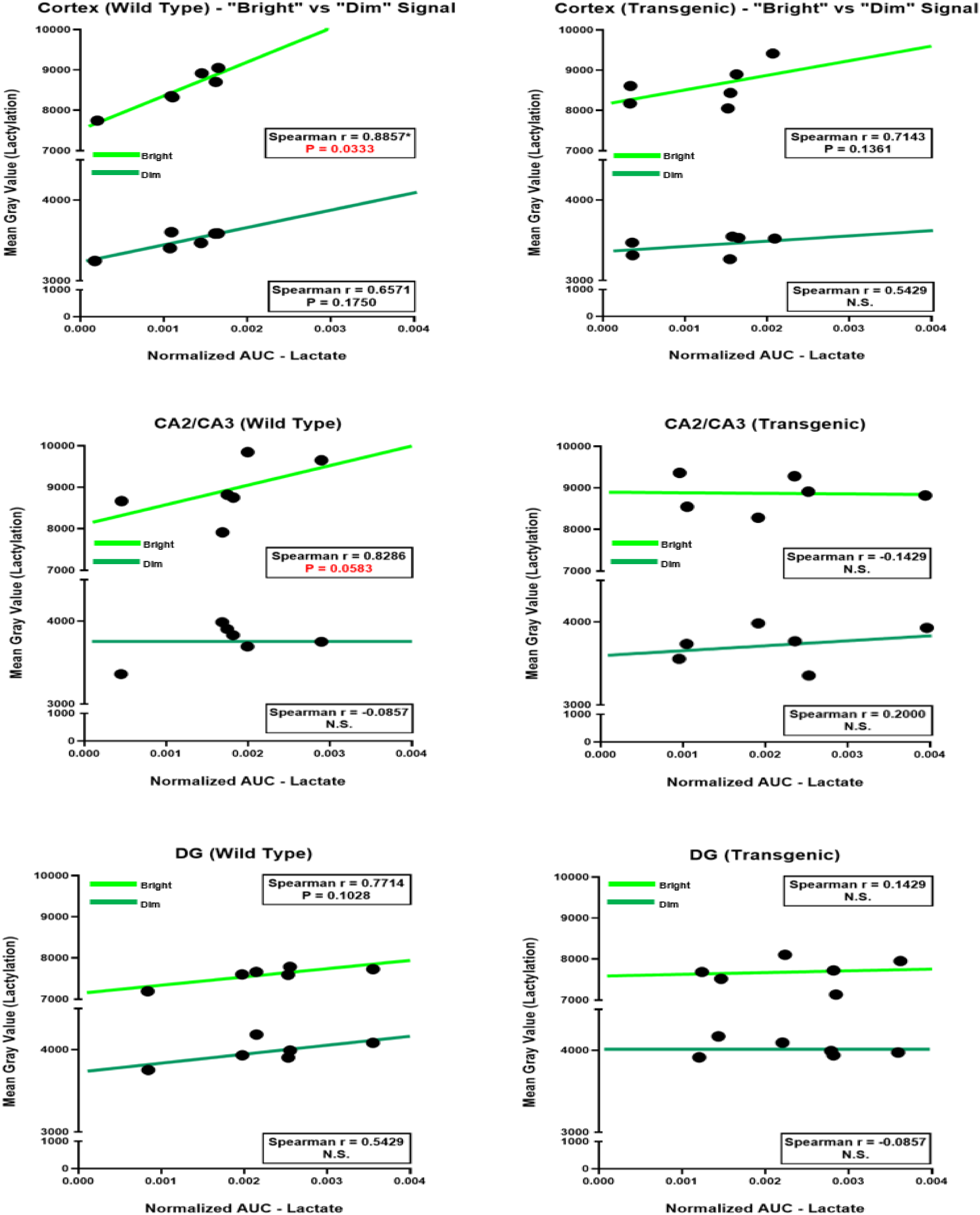
Correlational analysis of relative lactate levels determined from MALDI-TOF IMS data compared to mean gray values quantified through IF. A positive correlation was observed in the cortex, CA2/CA3, and DG ROIs of wild type mice at the putative non-neuronal, “Bright” intensity level (neon green line of best fit), but the association was not present in the cortex, the CA2/CA3 and DG subregions of transgenic mice. No significant difference was observed at the “Dim” signal level (dark green line of best fit) in both the cortex and DG (n=6).

Representative heatmaps were created to compare and contrast lactylation based on immunofluorescent staining, with lactate levels acquired by MALDI IMS, in female mouse brain tissue **(Figure S6)**. In general, areas with increased lactate levels matched to areas of higher lactylation signal **(Figure S6B & C)**. However, closer examination of the IF-based heat maps between transgenic versus wild type tissue found subtle differences in uniformity of staining whereby densely stained neuron-rich regions within the cerebellum and hippocampus were uniform and more intense in the wild type tissue but non-uniform or less intense within transgenic tissue **(Figure S6B)**. Lactylation found in the cortex and thalamus appeared brighter and distinct in transgenic tissue compared to wild type tissue. There also appeared to be less lactylation signal in the *stratum radiatum* layer of the CA1 subregion, but it was more pronounced in wild type compared to transgenic tissue.

## Discussion

A primary objective of this study was to establish a MALDI-TOF IMS workflow capable of detecting and spatially mapping lactate in mouse brain tissue. Few studies have applied MALDI-TOF IMS to central nervous system energy metabolism, and lactate is rarely examined using this approach. By employing 1,5-diaminonaphthalene (DAN) as the matrix, we demonstrate that lactate, along with related metabolic intermediates, can be detected and spatially resolved across discrete brain regions, enabling a region– and sex-specific analysis of cerebral metabolism.

### Age– and sex-dependent metabolic remodeling of the brain

Using this approach, we identified strong effects of age and sex on the distribution of lactate, glutamate, pyruvate, and citrate, particularly within hippocampal subregions and the thalamus **(Figures 1-4, 7B & C)**. Lactate and glutamate levels were highest in young mice and declined with age, with females consistently maintaining higher levels than males. In contrast, pyruvate levels were lowest in young animals and increased with age, most prominently in females, while citrate exhibited greater inter-cohort variability, with generally higher levels in males at 6 months and females at 18 months.

Together, these patterns suggest a progressive, age-related reduction in glycolytic activity across brain regions, accompanied by a relative shift toward oxidative metabolism, particularly within white matter. Directional ratio analyses supported this interpretation, revealing a robust increase in pyruvate-to-lactate ratios with age and region-specific changes in pyruvate-to-citrate ratios consistent with altered metabolic flux **(Figures 5, 6, 7B)**.

### Lactate declines with age but remains elevated in females

Previous work using *in vivo* approaches has reported elevated lactate in APP/PS1 mice relative to wild type controls at midlife [36]. In contrast, we observed a general age-related decline in lactate levels in both genotypes, with sustained elevation in females relative to males **(Figures 1, 7B & C)**. These differences likely reflect methodological distinctions. Prior studies relied on *in vivo* ^1^H magnetic resonance spectroscopy (^1^H MRS) or microdialysis, whereas MALDI-TOF IMS analyzes post-mortem tissue and allows finer regional resolution [36,44,51,58–60]. Moreover, lactate levels are influenced by circadian state and glymphatic clearance, which is enhanced during sleep [61–63]. Since animals were euthanized during the light phase and tissues were perfused prior to freezing, extracellular lactate may have been partially cleared, potentially contributing to lower absolute levels.

Notably, despite these factors, female mice, particularly transgenic females, retained higher intracellular lactate levels than males. This persistence suggests impaired lactate clearance or altered intracellular handling in females, a phenomenon that may be exacerbated in the presence of amyloid pathology. Additionally, our regionally targeted analysis excluded hippocampal compartments, such as the *subiculum* and *stratum lacunosum-moleculare*, which may contribute substantially to lactate pools measured in whole-tissue assays, further underscoring the importance of spatial resolution when interpreting metabolic data.

### Aging preferentially alters glycolysis rather than OXPHOS

Across both genotypes, age-related changes in glycolysis were more pronounced than changes in oxidative phosphorylation **(Figures 5 & 6)**. In grey matter regions, baseline OXPHOS activity was largely maintained with age, whereas glycolytic activity declined substantially, particularly in females. MALDI IMS images supported this interpretation, showing reduced lactate signal with age alongside region-specific increases in citrate signal. In males, a low but persistent level of glycolysis was maintained across the brain, with elevated OXPHOS activity evident in the hippocampus of aged transgenic mice.

These findings align with human positron emission tomography (PET) studies demonstrating that cognitive decline correlates strongly with reduced glucose consumption and aerobic glycolysis rather than changes in oxygen utilization [64–66]. Thus, the global decline in lactate observed here likely reflects a fundamental shift in cerebral glycolysis with aging, rather than a uniform reduction in mitochondrial metabolism.

### Sex as a dominant determinant of cerebral metabolism

In this study, the dominant influence of sex on cerebral metabolite levels emerged. Lactate and glutamate were consistently elevated in females at both ages, while pyruvate levels diverged between sexes with age **(Figures 1-3)**. These findings highlight an inherent sex-specific metabolic phenotype that persists under physiological aging and may shape vulnerability to neurodegenerative stress.

Despite growing recognition of sex differences in Alzheimer’s disease risk and progression, many metabolic studies either exclude females or fail to perform sex-stratified analyses [10,42,63,65,67–69]. Our data underscore the importance of explicitly incorporating sex as a biological variable when examining brain energy metabolism and its dysregulation in aging and disease.

### Sex-specific regulation of histone lactylation

Elevated lactate levels in females were paralleled by increased histone lactylation in aged female mice, particularly within the cortex and CA2/CA3 hippocampal subregion **(Data not shown)**. In transgenic females, lactylation was consistently enriched in regions proximal to amyloid plaques, a pattern that was especially pronounced in putative microglial populations **(Figure 8)**. These spatial patterns closely resemble prior observations that microglial lactylation increases in response to amyloid stress, although previous studies did not assess sex as a modifying factor [42,70].

The elevated pyruvate, citrate, and glutamate levels observed in aged females further suggest broader metabolic reprogramming that may support additional histone modifications via acetyl-CoA and α-ketoglutarate availability **(Figures 2-4)** [42,58,71,72]. Together, these data support a model in which sex-specific metabolic states influence epigenetic regulation of neuroinflammatory responses, with females exhibiting heightened sensitivity to amyloid-associated stress.

### Divergent coupling of lactate and lactylation in wild type versus transgenic mice

By integrating MALDI IMS with immunofluorescence, we examined the relationship between lactate abundance and histone lactylation. In wild type mice, increasing lactate levels correlated strongly with increased lactylation, particularly in regions and cell types consistent with microglia **(Figure 9)** [73,74]. Surprisingly, this relationship was weakened or absent in transgenic mice, despite evidence of elevated lactylation near plaques **(Figures 8, S4, S5B)**.

This decoupling suggests that amyloid pathology alters the relationship between metabolic state and epigenetic regulation, potentially through chronic microglial activation, altered substrate utilization, or impaired metabolic flexibility [42,58,70–72]. The pronounced sex dependence of this effect further indicates that females may engage distinct metabolic–epigenetic pathways in response to amyloid stress.

### Limitations and future directions

This study is limited by the absence of behavioral assessments, precluding direct correlations between metabolic changes and cognitive status [64]. Survivorship bias may also influence findings at 18 months [75]. Future studies should incorporate intermediate ages, behavioral testing, and longitudinal designs. Additionally, cell-type specificity was inferred from staining intensity rather than confirmed using markers for microglia, astrocytes, or oligodendrocytes; an important direction for future work, particularly given the metabolic demands of myelination and astrocytic reactivity in AD [64,73,75–81].

### Conclusions

This study establishes MALDI-TOF IMS as a powerful tool for spatially resolved analysis of brain metabolism and reveals sex-specific metabolic and epigenetic signatures associated with aging and amyloid pathology. Despite technical challenges inherent to lactate detection, we demonstrate that female mice maintain elevated cerebral lactate and histone lactylation with age, particularly in regions vulnerable to AD. These findings support a model in which impaired lactate clearance and microglial metabolic reprogramming may contribute to sex-dependent vulnerability in neurodegeneration and highlight histone lactylation as a promising mechanistic link between metabolism and neuroinflammation.

## Acknowledgements

We would like to thank Dr. Chaochao Chen and Dr. Ken Yeung (Department of Biochemistry, Western University) for their technical assistance and advice.

