## Supplemental Figures for "MALDI-TOF imaging mass spectrometry demonstrates sex- and age-dependent spatial changes in brain energy metabolism in response to amyloid stress using a mouse model of Alzheimer’s Disease"

[a + b] Corresponding authors

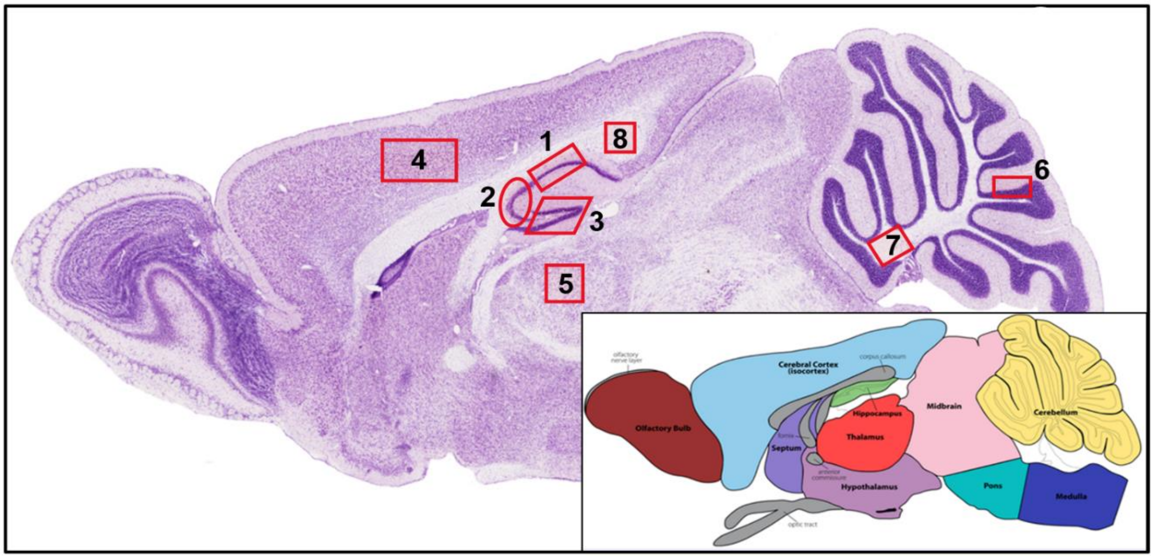


**Figure S1: Hematoxylin and eosin (H&E) stain of the mouse brain (sagittal section)**

The sagittal section displayed highlights all 9 ROIs that were selected for MALDI IMS analysis: (**1**) CA1 subregion of the hippocampus, (**2**) CA2/CA3 subregion of the hippocampus, (**3**) DG, also part of the hippocampus, (**4**) cortex, (**5**) thalamus, (**6**) cerebellum, (**7**) white matter of the cerebellum, (**8**) corpus callosum, and the entire brain section. Only regions 1 through 4 were further analyzed for lactylation post immunofluorescent staining [Images obtained from the Gene Expression Nervous System Atlas (GENSAT) Project [57]].

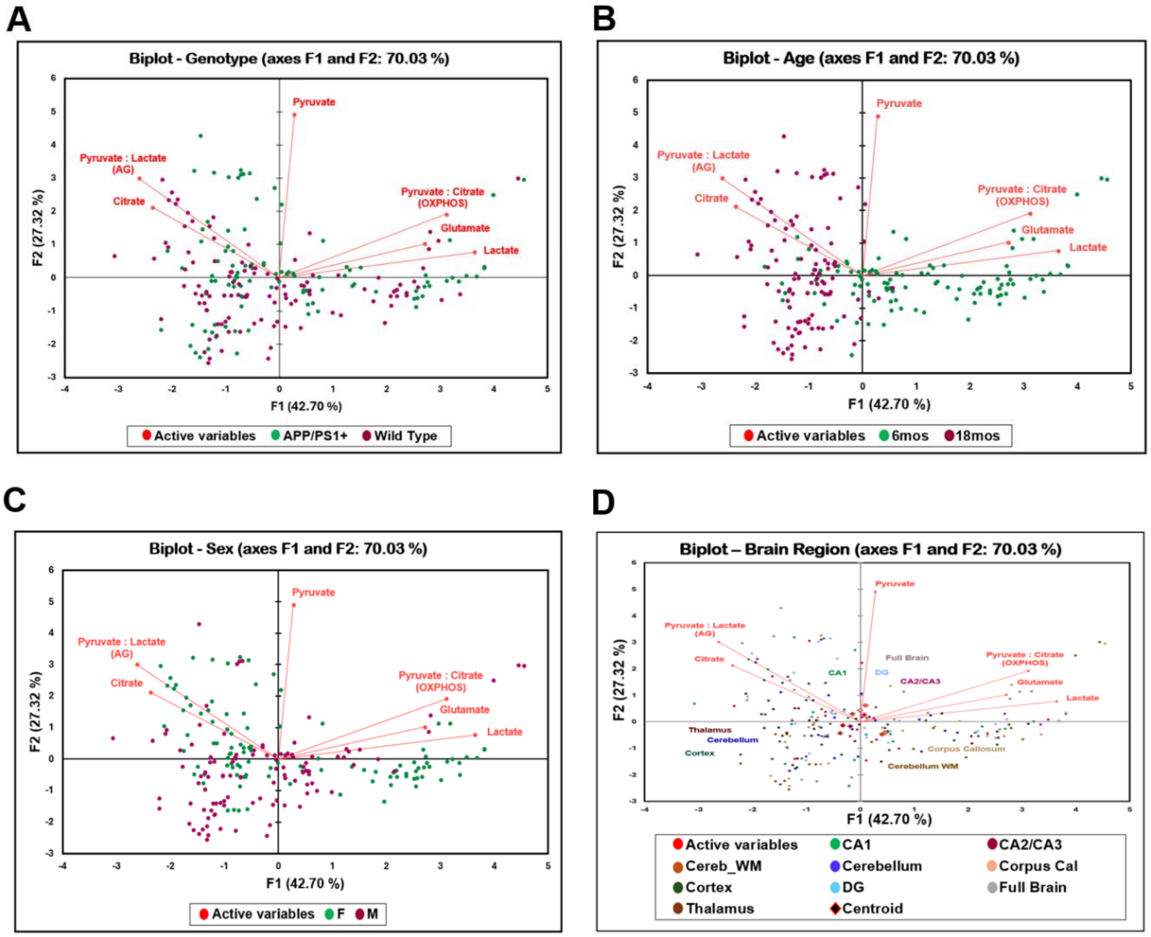


**Figure S2: Principal component analysis of cerebral metabolite levels and metabolic flux based on genotype, age, sex, and brain region**

Levels of lactate, glutamate, and pyruvate as well as the directional ratio of pyruvate to citrate were predominantly associated with the younger 6-month-old mice within brain regions that included the CA2/CA3 and DG subregions of the hippocampus along with the full brain (**A-D, top right quadrants**). Citrate and the directional ratio of pyruvate to lactate were predominantly associated with the aged 18-month-old mice within the CA1 subregion of the hippocampus (**A-D, top left quadrants**). There was no clear pattern of association present based on genotype.

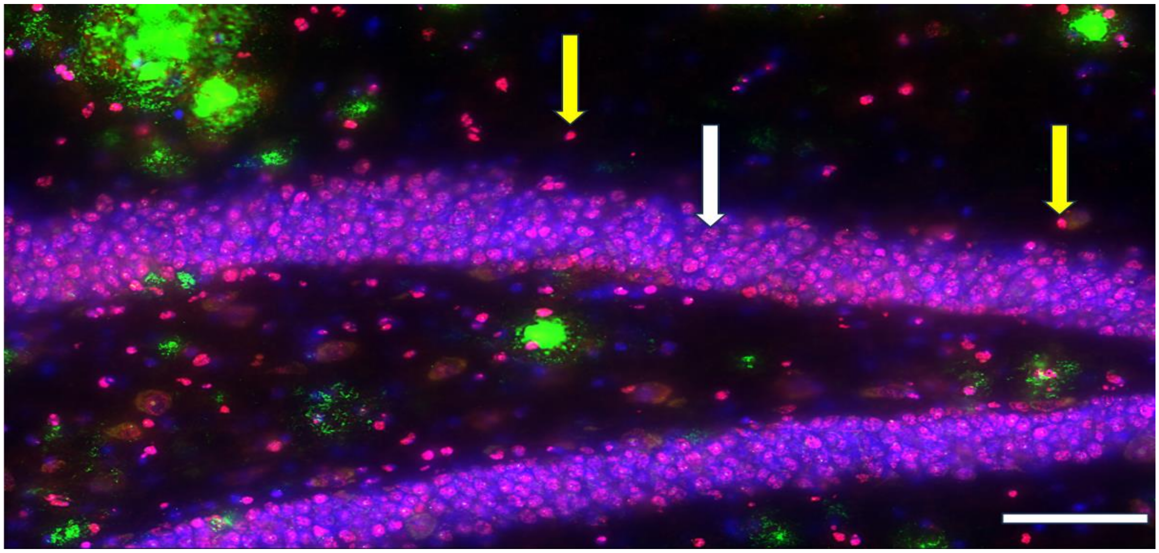

**Figure S3: Differential histone lactylation staining within the mouse dentate gyrus**

Eighteen-month-old transgenic AD mouse brain tissue was dual stained with amyloid- and lactylation-specific antibodies. Amyloid-containing plaques and lactylated proteins were highlighted in green and red, respectively, while nuclei were counterstained with DAPI, highlighted in blue. Brighter or more intense lactylation signal was predominantly localized to small, nucleated cells typical of microglia (yellow arrows) whereas larger nuclei in the neuron dense granular cell layer of the dentate gyrus revealed a dimmer or less intense lactylation signal (white arrow). Deconvolved image, 60x magnification, scale bar represents 100 µm.

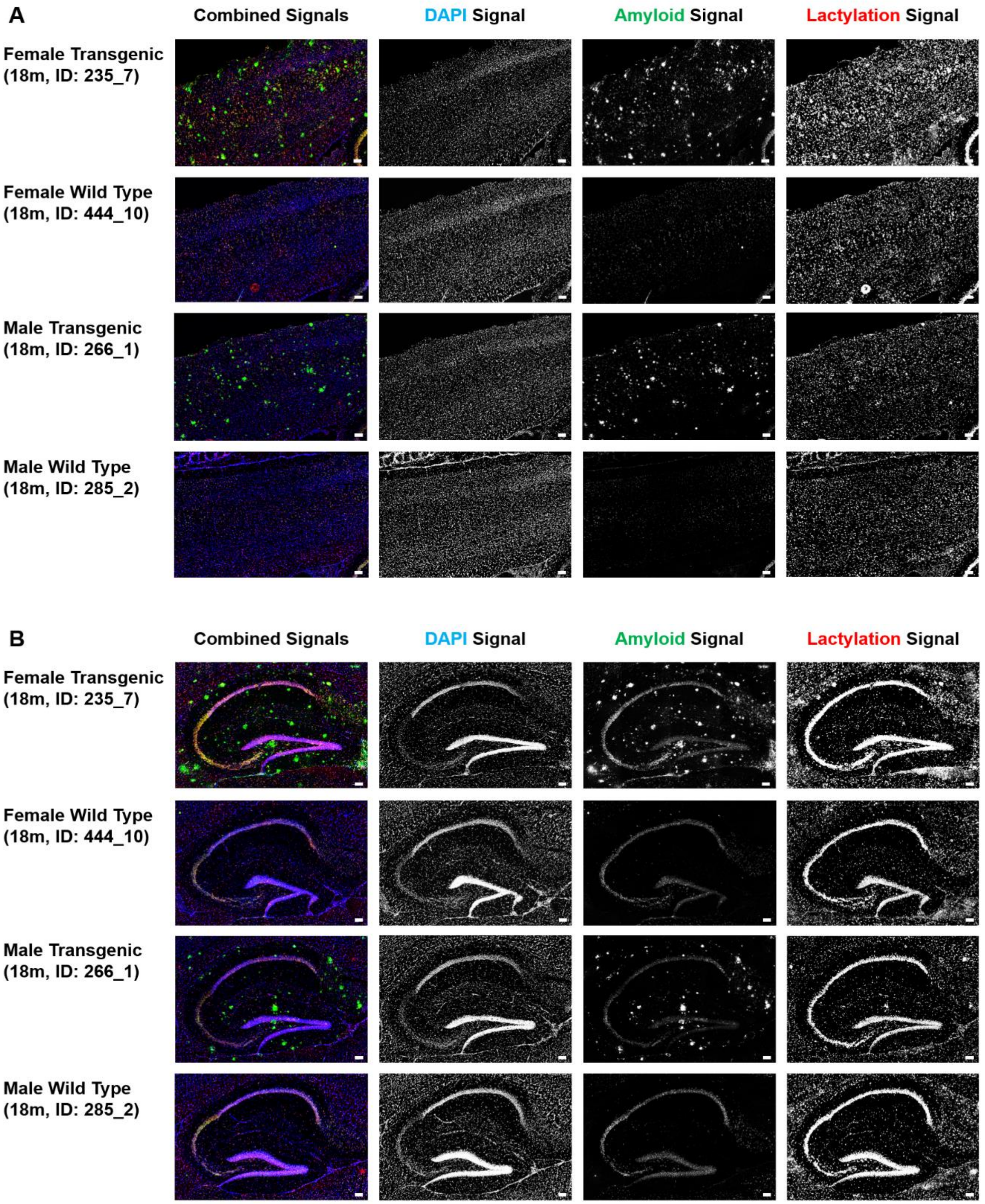


**Figure S4: Immunofluorescent staining of amyloid and lactylated proteins in the cortex and hippocampus of 18-month-old transgenic AD mice**

Eighteen-month-old brain tissues from transgenic male and female mice were analyzed by immunofluorescence microscopy using dual staining for amyloid (green) and protein lactylation (red) in the cortex (**A**) and the hippocampus (**B**). From the 4 ROIs selected for analysis, the amyloid burden was greatest in the cortex and in the DG subregion of the hippocampus. Scale bars represent 100 μm.

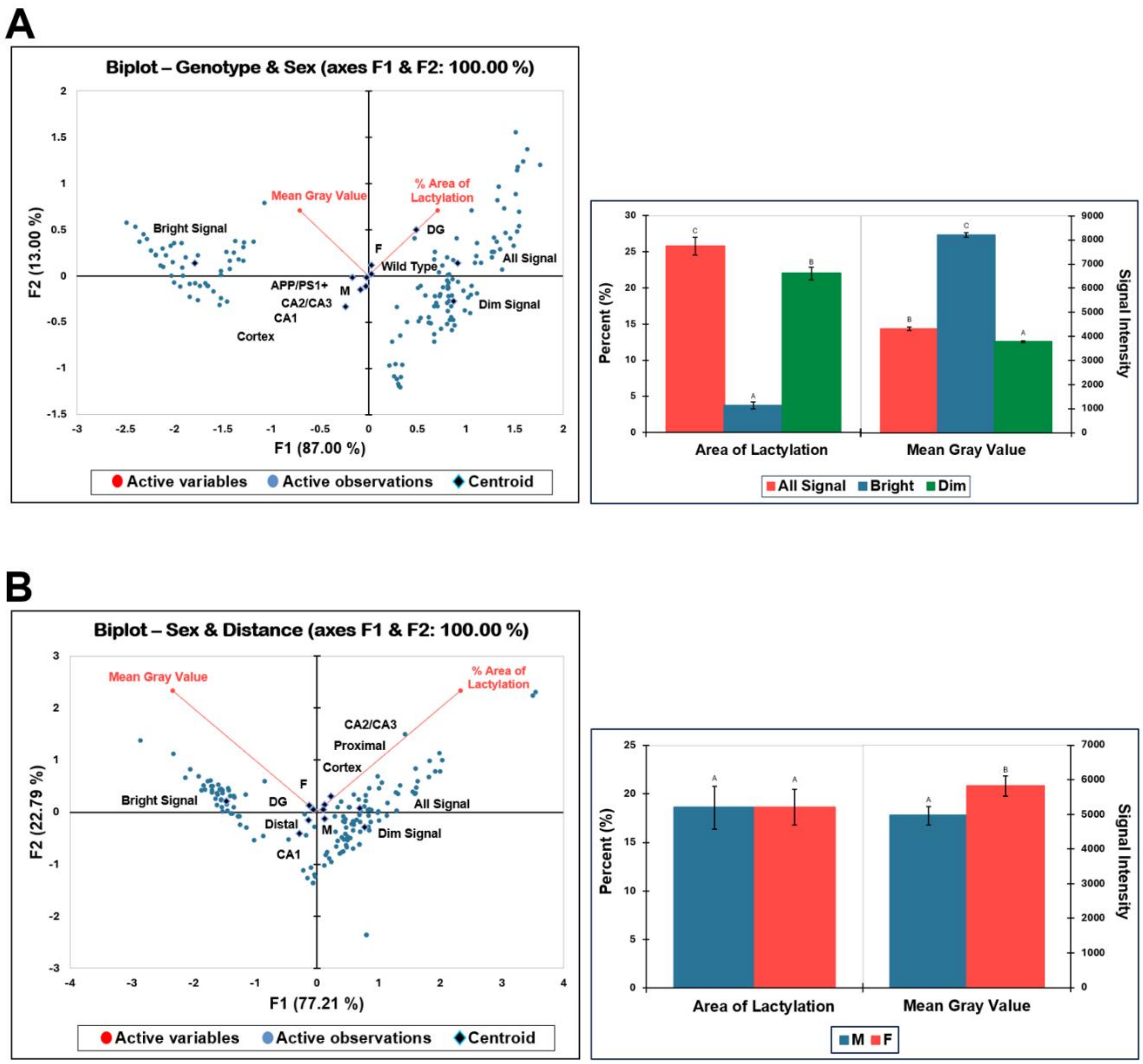


**Figure S5: Principal component analysis of histone lactylation in mouse brain tissue**

The ordination of protein lactylation in tissues from both transgenic and wild type mice was determined for 4 anatomical brain regions: the cortex, CA1, CA2/CA3, and DG. The percent area of all lactylation signal associated with wild type females within the DG (**A, top right quadrant**). Alternately, the mean gray value was associated with the bright lactylation signal (**A**, **top left quadrant**). One-way ANOVA also confirmed the 3 significantly different intensity ranges of lactylation signal (n=48) (**A**). However, in transgenic mice brain tissues, the percent area of all lactylation signal was associated with regions proximal to plaques within the cortex and CA2/CA3 (**B, top right quadrant**) while the mean gray value was specifically associated with the bright lactylation signal found in the DG of female mice (**B, top left quadrant**). Furthermore, the mean gray values were significantly elevated in aged transgenic female mice compared to aged transgenic male mice (n=72) (**B**).

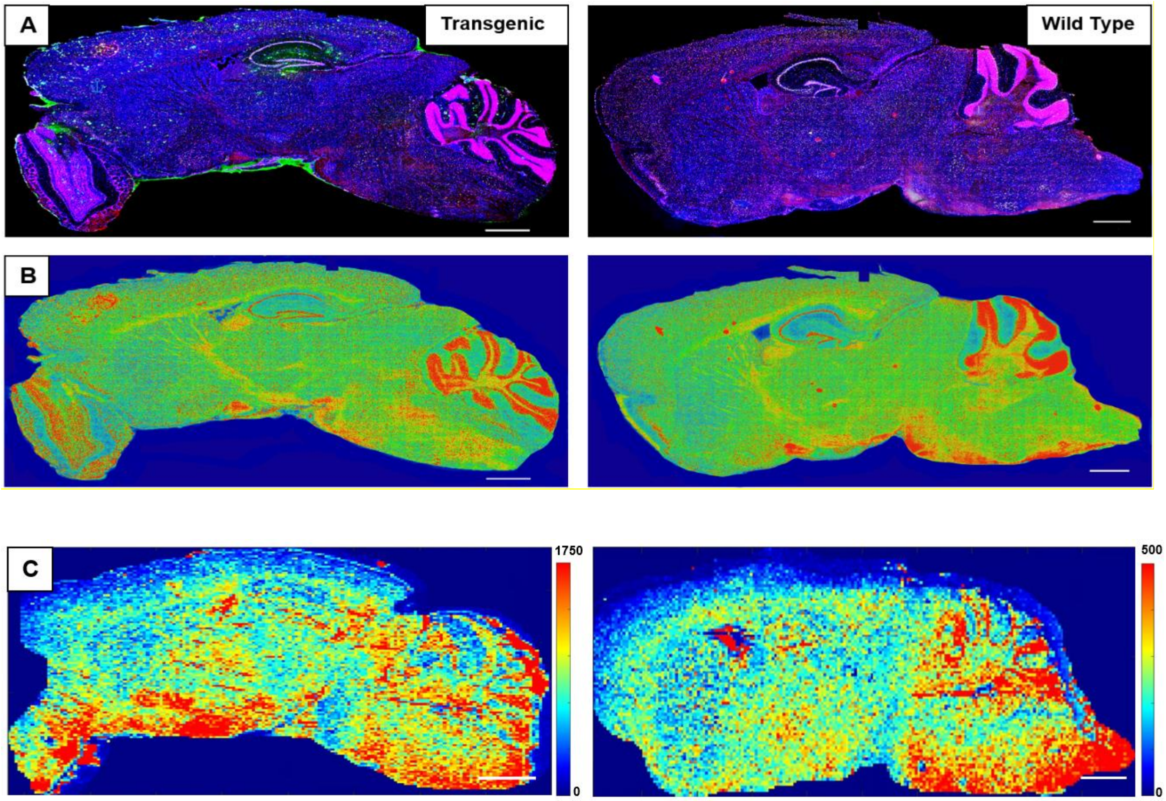


**Figure S6: Whole brain images of lactylation signal captured by immunofluorescence microscopy compared to lactate levels detected with MALDI-TOF IMS**

Large image tiles were stitched to create whole brain scans of sagittal mouse sections following immunofluorescence staining using an anti-lysine-lactylation antibody (red) with a DAPI nuclear counterstain (blue) and imaged with a Nikon inverted Ti2E deconvolution microscope, at 20x magnification (panel **A**). A representative female transgenic AD mouse brain is shown in the left column compared to a representative female wild type mouse brain contained in the right column. The fluorescent signal from the anti-lysine-lactylation antibody was converted into a heatmap using NIS-Elements software to represent lactylation signal (panel **B**). An additional heatmap highlighting lactate signal detected at m/z 89 by MALDI-TOF IMS was also generated through MSiReader for comparison (panel **C**); the highest level of signal is represented in red. In general, areas with increased lactate levels matched to areas of higher lactylation signal. However, closer examination of the IF-based heat map (panel **B**) found subtle differences in the uniformity of staining possibly related cell-type specific variation in lactylation signal when transgenic to wild type images were compared. The scale bars for IF- and IMS-based images represent 1000 µm and 20 mm, respectively; lateral resolution of 70 µm for IMS-based images.
